# NLCD: A method to discover nonlinear causal relations among genes

**DOI:** 10.64898/2026.03.20.713150

**Authors:** Aravind Easwar, Manikandan Narayanan

## Abstract

Distinguishing correlation from causation is a fundamental challenge in many scientific fields, including biology, especially when interventions like randomized controlled trials are infeasible and only observational data are available. Methods based on statistical tests of conditional independence within the Mendelian Randomization framework can detect causality between two observed variables that are each associated with a third instrumental variable. However, these methods for detecting causal relationships between traits (e.g., two gene expression or clinical traits associated with a genetic variant, all observed in the same population) often assume a linear relationship, thereby hindering the discovery of causal gene networks from genomics data.We have developed NLCD, a method for NonLinear Causal Discovery from genomics data based on nonlinear regression modeling and conditional feature importance scoring. NLCD uses these techniques to extend the statistical tests in an existing linear causal discovery method called the Causal Inference Test (CIT). We benchmarked NLCD against current state-of-the-art methods: CIT, Findr, and MRPC. On simulated datasets, NLCD performs comparably to most methods in detecting linear relations (Average AUPRC (Area Under the Precision-Recall Curve) of NLCD=0.94, CIT=0.94, Findr=0.94, and MRPC=0.99), and outperforms them in detecting nonlinear (sine and sawtooth type) relations between two genes (Average AUPRC of NLCD=0.76, CIT=0.60, Findr=0.56, and MRPC=0.73). When tested on a nonlinear subset of a yeast genomic dataset to recover known causal relations involving transcription factors, NLCD and CIT performed comparable to each other and slightly better than Findr and MRPC (Average AUPRC of NLCD=0.82, CIT=0.81, Findr=0.71, and MRPC=0.54). On application to a human genomic dataset, NLCD revealed active causal gene pairs (*IRF1* → *PSME1* and *HLA-C* → *HLA-T*) in the muscle tissue, and clarified the promises and challenges in discovering causal gene networks in tissues under *in vivo* human settings.

**Availability:** The code implementing our method is available at: https://github.com/BIRDSgroup/NLCD.

## 1. Introduction

Discovering causal relationships among a set of variables that constitute a complex system has been an active area of research in many fields, including bioinformatics, due to its many applications. For instance, distinguishing genes that cause a disease from those that are merely affected by the disease is important for finding targets for new drugs/therapies vs. diagnostic biomarkers for the disease. But an observed correlation between two traits such as gene expression traits does not imply causation, as it could also result from a confounder variable affecting both traits (Pearl, 2009).

A gold standard solution for testing causality in the presence of confounding is a Randomized Controlled Trial (RCT), but RCTs are not always feasible and present additional challenges in an *in vivo* human setting (Klungel et al., 2004). An alternative solution for bivariate causal discovery is offered by Mendelian Randomization (MR) methods. MR-like methods that discover a causal relationship between two gene trait variables using a genetic variant associated with both genes in a population include CIT (Millstein et al., 2009) (and its precursor (Schadt et al., 2005)), Findr (Wang and Michoel, 2017), Trigger (Chen et al., 2007), and MRPC (Badsha and Fu, 2019; Badsha et al., 2021; Kvamme et al., 2025). These methods utilize the associated genetic variant (single nucleotide polymorphism or SNP) as an instrumental variable to control for confounding between the observed genes using only observational data.

Despite research spanning two decades on MR methods for gene-gene causal discovery mentioned above, a limitation of most methods is their assumption of a linear relationship between the two genes or clinical traits being tested. Many trait pairs could significantly deviate from linearity (e.g., 84% of all tested pairs in the UK BioBank data (Sulc et al., 2022)), and periodically changing genes (according to the circadian rhythm) could also exhibit nonlinear relationships (Sukumaran et al., 2010). A few studies have tried to incorporate nonlinearity (Sulc et al., 2022; Burgess et al., 2014; Arvanitis et al., 2021), but they assume a piecewise linear or polynomial relation between the traits. Hence, there is a need for a more general nonlinear causal discovery method.

This work advances causal discovery research by proposing a new method NLCD for NonLinear Causal Discovery. NLCD employs conditional importance scoring of features in a general nonlinear regression model, in order to generalize a key conditional independence test of causality from linear to nonlinear settings. Specifically, NLCD extends a linear causal discovery method, CIT, by conducting a series of permutation-based statistical tests of causality in the nonlinear setting. The maximum p-value from these tests helps decide if data on a given triplet (two gene trait variables and a third instrumental variable) supports causality vs. independence between the traits. NLCD can flexibly work with any nonlinear regression model (including Kernel Ridge Regression, Support Vector Regression, or Artificial Neural Network models).

In benchmarks comprising simulated triplets, NLCD showed better performance in identifying nonlinear causal relations (for example, Average AUPRC was 0.72/0.57/0.51/0.71 in the order for NLCD/CIT/Findr/MRPC on a nonlinear sawtooth-type dataset). When benchmarked on a nonlinear subset of a yeast genomic dataset, NLCD’s average performance in recovering known causal relations involving transcription factors was comparable to CIT and better than Findr and MRPC. Application of NLCD to a human multi-tissue genomic dataset (called the Genotype-Tissue Expression or GTEx dataset) revealed a few causal gene pairs that are active in the muscle tissue, and highlighted the difficulty and hence the care that should be taken in filtering the discovered causal relations, as well as advantages of a data-driven approach to discover *in vivo* human causal gene networks.

## 2. Methods

### 2.1 Triplet-based causal discovery problem

We are interested in the problem of testing whether a causal relationship exists between two trait variables *A* and *B* (e.g., two gene expression traits focused in this work; clinical traits can also be considered), when there exists a third instrumental variable *L* (e.g., a SNP capturing the genetic variation occurring at a nucleotide in the genome). Together, these three variables are called a triplet (see Fig. 1a).

**Fig. 1.**
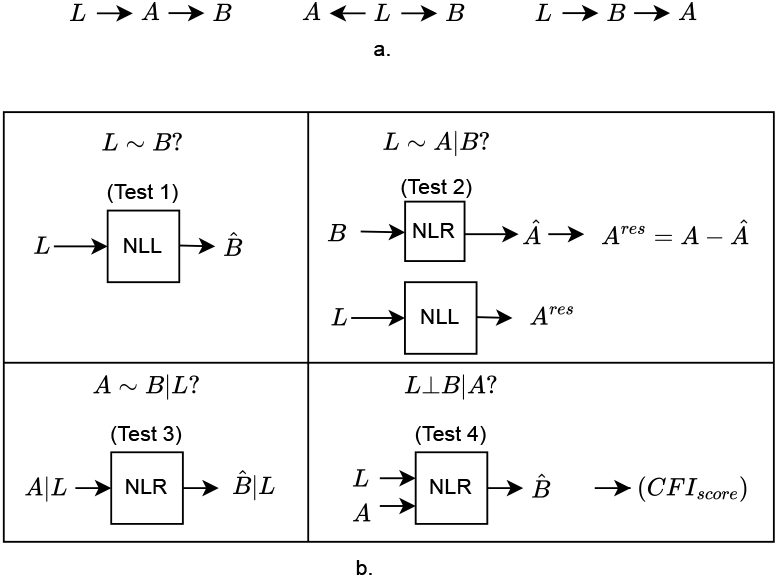
Schematic of our NLCD method. a. Given observational data on a genotype variable *L* and two gene expression variables *A* and *B* (measured across the same set of individuals), our method NLCD classifies the triplet (*L, A, B*) as following a causal (left), reactive (right), or independent (middle) model. b. To test if a triplet is causal, NLCD conducts four statistical tests: an association test (Test 1), two conditional association tests (Tests 2-3), and a key conditional independence test (Test 4). NLCD’s main contributions include the use of NonLinear Regression (NLR) modeling to perform three relevant tests and the development of a Conditional Feature Importance (CFI) score to implement Test 4. NLL stands for the Negative Log Likelihood test statistic.

#### Input

The triplet (*L, A, B*) can be thought of as three random variables that are observed in a set of *n* individuals, with the individuals randomly sampled from a population of interest. This observational (also referred to sometimes as observed or training) data on the triplet forms the input, and denoted as *D*_*T*_ = {(*L*_1_, *A*_1_, *B*_1_), (*L*_2_, *A*_2_, *B*_2_), …, (*L*_*t*_, *A*_*t*_, *B*_*t*_), …, (*L*_*n*_, *A*_*n*_, *B*_*n*_)} (see Table S1). Note that *L* is a discrete random variable representing a genetic factor (e.g., a SNP), and can have values 0 or 1 for a haploid individual (e.g., yeast segregant) or values 0, 1 or 2 for a diploid individual (e.g., human). *A* and *B* are continuous random variables representing a pair of gene expression or any other traits.

#### Output

The expected output is to call that the triplet follows one of the three models, causal, reactive, or independent (see Fig. 1a), based on the input observational data on the triplet. Output includes p-values associated with the call. Output can also be “No call” if evidence is inconsistent or inconclusive.

#### MR-like assumptions/requirements

Any method that solves the above causal discovery problem need to make certain assumptions. We are interested in only a class of methods that make the key assumption that *L* is “randomized”, i.e., if each individual’s *L* value is randomly set (to one of its possible alleles) before and independent of the individual’s *A* and *B* (gene expression) values. As explained in Supplementary Section 1.1 Background on underlying theorem and model, this assumption is one of the standard assumptions in MR as it is supported by randomness in biological processes such as reproductive cell formation; and enables development of methods that use *L* as a valid instrument or “causal anchor” to infer the causal relation between *A* and *B* based on an established causal equivalence theorem (Chen et al., 2007; Millstein et al., 2009).

For convenience, certain methods make a few additional assumptions about the input triplet that can be easily verified using the triplet’s data. We can assume that the two genes in the triplet are correlated (i.e., *A* ∼ *B*), since only correlated genes need be tested for causation. For *L* to be an instrument, we can also assume that *L* is correlated to both *A* and *B*. If the allelic variation at *L* is associated with the expression variation of a gene *G* across individuals, then this correlated (*L* ∼ *G*) pair is called an expression Quantitative Trait Loci (eQTL) relation, with *L* being the eQTL SNP and *G* being the eQTL gene in the pair. Methods can filter triplets to focus only on triplets where *L* is an eQTL SNP for both *A* and *B*.

### 2.2 Our NLCD method

Our NLCD method solves the triplet-based causal discovery problem by generalizing existing approaches, which typically assume a linear relationship between *A* and *B*, to the nonlinear setting. NLCD’s key contributions are to use a NonLinear Regression (NLR) model to capture the nonlinear causal relation between *A* and *B*, and to develop conditional feature importance (CFI) and related measures to properly realize four statistical tests that establish causality as per the causal equivalence theorem. We describe these contributions after providing some preliminaries. Note that NLCD can also discover linear causal links, as linearity is a special case encompassed by the broader nonlinear relations modeled by NLR.

#### 2.2.1 Preliminaries

To clarify the context for our NLR models and statistical tests of the causal equivalence theorem (Fig. 1), we explicitly provide the joint distribution of the three models of interest. For a random variable *X*, let *X* ∼ 𝒩 (*µ, σ*^2^) denote that *X* is sampled from or follows a univariate Gaussian distribution with mean *µ* and variance *σ*^2^, and 𝒩 (*x*; *µ, σ*^2^) denote the probability density function of this Gaussian random variable *X* evaluated at *x* (i.e., 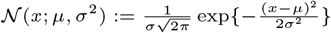).

Following standard probabilistic graphical model framework (Koller and Friedman, 2009), the joint distribution for the causal model *L* → *A* → *B* is given by *P* (*A, B, L*) = *P* (*L*) *P* (*A* | *L*) *P* (*B* | *A*), reactive model is similarly factorized, and (conditional) independence model *A* ← *L* → *B* is given by *P* (*A, B, L*) = *P* (*L*) *P* (*A* | *L*) *P* (*B* | *L*). Here *P* denotes the appropriate probability mass or density function. For instance, the prior *P* (*L*) is the probability mass function of a Bernoulli distribution (for haploids) or multinoulli/categorical distribution (for diploids). The conditional 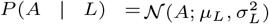, where the mean and variance parameters depend on the value of *L. P* (*B* | *L*) is similarly defined but using its own set of parameter values (denoted 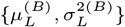, or simply 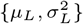 by abusing notation when the context is clear about gene *B*). To complete the specification of the joint distribution, we need to specify the conditional *B* | *A* as described next.

#### 2.2.2 NonLinear Regression (NLR) modeling

NLCD captures nonlinear relationship via the conditional distribution (*B* | *A*) ∼ 𝒩 (*f* (*A*), *σ*^2^), where *f* (.) is a nonlinear function of its argument. While testing for conditional independence, we also need another conditional distribution, which we model as (*B* | *L, A*) ∼ 𝒩 (*g*(*L, A*), *σ*^′2^), where *g*(., .) is another nonlinear function of its arguments.

To learn these nonlinear functions *f* (.) and *g*(.) from data, we employ NLR. Simple NLR models suffice due to *L, A*, and *B* each being an one-dimensional random variable. We specifically use the following models from the scikit-learn library (Pedregosa et al., 2011): Artificial Neural Network (ANN) with one hidden layer, Support Vector Regression (SVR) with radial basis function (RBF) as the kernel, and Kernel Ridge Regression (KRR) with RBF as the kernel. The relevant scikit-learn functions were called as detailed in Supplementary Section 1.2 Specifications of the nonlinear models used. Note that the learnt *f* (.) and *g*(.) may be linear when the data supports it, as nonlinear functions include linear functions as a special case.

#### 2.2.3. Conditional Feature Importance (CFI) score

To realize a conditional independence test that is needed to discover causality, we need a corresponding statistic that captures the strength of the relation between two variables (*L* and *B* in our case), conditional on a third variable (*A* in our case). We propose a CFI score for this purpose; it is inspired by the conditional predictive impact measure from an earlier work (Watson and Wright, 2021).

Consider an NLR model to predict *B* from *L* and *A* (denoted *B* | *L, A*), and let it be trained on the triplet’s data 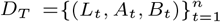. Let 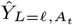 be the prediction of this model when *L* = *ℓ* and *A* = *A*_*t*_. Let

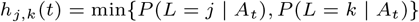

where *j, k* ∈ {0, 1} for haploids and *j, k* ∈ {0, 1, 2} for diploids (and the relevant probabilities calculated using Bayes’ theorem as shown in Supplementary Section 1.3 Calculation of probabilities for CFI score). Our proposed CFI score for a dataset involving haploids is then:

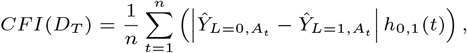

and diploids is:

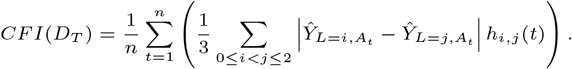

We now provide rationale for the haploid CFI score, as the diploid CFI’s rationale is similar. The first term in the summand of haploid CFI quantifies the importance of feature *L* conditional on feature *A* = *A*_*t*_ in an NLR model for predicting *B* (and thus helps test *L* ⊥ *B* | *A*). But this term is not trustable if 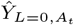 or 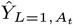 is not trustable – which is the case when the NLR model has never seen datapoints in the vicinity of (*L* = 0, *A* = *A*_*t*_) or (*L* = 1, *A* = *A*_*t*_) during training. Therefore, the second term in haploid CFI’s summand, *h*_0,1_(*t*), gives more weight to *A*_*t*_ values for which both *L* = 0 and *L* = 1 are likely to be seen during training.

To provide further insight into CFI, we define a related “overlap score” for a dataset involving haploids as:

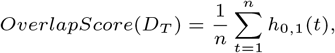

and diploids as:

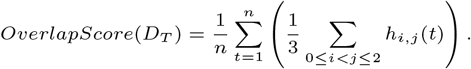

Intuitively, the haploid overlap score captures the overlap between the support of two gene expression distributions (*A* | *L* = 0 and *A* | *L* = 1) and CFI captures the difference in predictions among different values of *L* in the regions of overlap. Similar intuition also applies to the diploid overlap score and CFI.

#### 2.2.4. Realizing the four statistical tests of causality

As mentioned earlier, NLCD is based on an established causal equivalence theorem (Chen et al., 2007), where a chain of three statistical tests was used to test for causality (see Supplementary Section 1.1 Background on underlying theorem and model). We rely actually on a modified version of this theorem (Millstein et al., 2009), where the tests were slightly changed by conditioning on different variables and a fourth test was added to improve empirical performance (of a linear causal discovery method CIT). Our method NLCD extends the implementation of these four statistical tests (shown below and also in Fig. 1b) from the linear to nonlinear setting using the concepts of NLR modeling and CFI scoring:

(Test 1): *L* ∼ *B*

(Test 2): *L* ∼ *A* | *B*

(Test 3): *A* ∼ *B* | *L*

(Test 4): *L* ⊥ *B* | *A* (Conditional Independence Test)

NLCD calculates the empirical p-value for each test using permutation testing. In detail, we calculate the test statistic on the actual observed data and several permuted datasets (with permutation appropriately done to make the permuted data conform to the null model). The empirical p-value is then estimated using the number of permutations (*m*) in which the statistic is as extreme as or more extreme than the statistic calculated on the actual observed data, as follows. If *M* is the total number of permutations (default *M* = 500), the estimated empirical p-value is given by 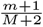. The pseudo counts of +1 and +2 in this formula handle cases where *m* = 0. We now describe the statistic and permutation scheme used by NLCD in its four tests.

- **Test 1**: *L* ∼ *B* This is a test of the association between *L* and *B*. Regardless of the underlying true model generating the data, we assume for this test that 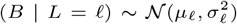. Let *n*_*ℓ*_ be the number of training datapoints with genotype value *ℓ*, i.e., *n*_*ℓ*_ := |{*t* ∈ {1, …, *n*} : *L*_*t*_ = *ℓ*}|. Then, we use the training data to compute the maximum likelihood estimates (MLEs) of (*B* | *L* = *ℓ*) as:

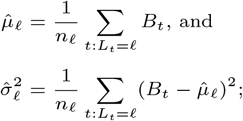

and use these MLEs computed for each possible value of *ℓ* to compute the Negative Log Likelihood (NLL) loss (i.e., observed test statistic) as:

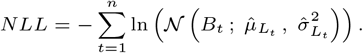

Next, we randomly permute the *B* trait (i.e., the vector 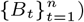 to break any association with *L*, and recompute the MLEs and NLL loss on this permuted dataset – this permutation-based NLL statistic is expected to be higher than the observed statistic if *L* is truly associated with *B*. Finally, repeating this process for *M* permutations and comparing the resulting statistics with the observed statistic as explained above yields the empirical p-value for this test.
- **Test 2**: *L* ∼ *A* | *B* We test the association of *L* and *A* given *B*. We use the training data to learn a NLR model to predict *A* from *B*, i.e., learn a nonlinear function *f* (.) such that *A* ≊ *f* (*B*). Then, for the *t*-th training datapoint, we have the prediction *Â*_*t*_ = *f* (*B*_*t*_), and residual 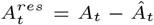. Let *A*^*res*^ denote the residual variable (and also the vector 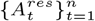). We finally test *L* ∼ *A*^*res*^ using the same association test described above (Test 1), but with *A*^*res*^ used in place of *B* to get a p-value for association between *L* and (*A* | *B*).
- **Test 3**: *A* ∼ *B* | *L* This test detects association between *A* and *B* in any strata of *L*. To do so, we obtain a p-value of association between *A* and *B* for each possible value of *L* as explained next, and take the minimum of the obtained p-values as the final p-value of Test 3. For any given value *ℓ* of *L* (with *ℓ* ∈ {0, 1} in case of haploid and *ℓ* ∈ {0, 1, 2} for diploid), we use the subset of training datapoints with *L*_*t*_ = *ℓ* to fit a NLR model to predict *B* from *A*, and take the associated mean square error (MSE) loss shown below as the observed test statistic:

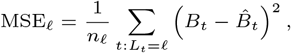

where 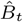 is the model prediction based on input *A*_*t*_. Next, as in Test 1, we independently and randomly permute the *B* trait but within the stratum defined by *L* = *ℓ* (i.e., the vector {*B*_*t*_ : *L*_*t*_ = *ℓ*}) to break any association with *A* within this strata, and refit the NLR model again within strata *ℓ* and recompute the MSE_*ℓ*_ loss (which is expected to be higher than the observed statistic if there is true association). Repeating this process *M* times and comparing the resulting permutation-based statistics with the observed statistic yields the empirical p-value of *A* ∼ *B* given *L* = *ℓ*.
- **Test 4 (Conditional independence test)**: *L* ⊥ *B* | *A* To test this conditional independence, we first fit an NLR model, with *L* and *A* as inputs and *B* as output, to the training data; and use the associated CFI score as the observed test statistic (see Sec. 2.2.3 Conditional Feature Importance (CFI) score on the derivation of CFI, and how CFI is a relevant test statistic quantifying the conditional dependence between *L* and *B* given *A*). Next, we seek a permutation scheme for shuffling the observed data to match a stringent null model—one that eliminates the conditional independence of interest while preserving all other inter-variable relationships as much as possible. One such scheme is a random permutation of *B* stratified by *L*. This permutation breaks any dependence between *B* and *A* within each stratum defined by *L*, thereby preserving any marginal dependence between *B* and *L*, and importantly enforcing the null model 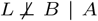 (Millstein et al., 2009). Note that the null model cannot be enforced if there is no marginal *B* ∼ *L* dependence in the observed data; however this is not a concern as Test 1 can filter out such triplets. The process to compute a permutation-based CFI comprises these steps: independently and randomly permute the values of *B* within each stratum defined by *L* to obtain the permuted vector *B*_stratperm_, fit an NLR model to the training data with *B* replaced by *B*_*stratperm*_, and compute the associated CFI (which is expected to be greater than the observed CFI if *B* is truly conditionally independent of *L* given *A*). As before, repeating this process *M* times and comparing the resulting statistics with the observed statistic yields the empirical p-value for Test 4.

#### Obtaining the final p-value

Following the intersection-union test framework (Cassella and Berger, 2002), we take the maximum of the p-values from the four tests above to get a single p-value. On this final p-value, we apply a suitable cutoff (say a significance level of 0.05) to decide if the triplet is causal (*L* → *A* → *B*) or not.

#### 2.2.5. Making the final call

If we know *a priori* that *A* is the potential regulator/exposure variable and *B* is the potential target/outcome variable, then we need to apply NLCD tests only in the forward direction *L* → *A* → *B* and compare it to a cutoff (say 0.05) as discussed above to call whether the triplet is causal or not. We make NLCD calls in this fashion when analyzing simulated and yeast benchmark datasets.

If the causal direction is not known *a priori* like in the human dataset, then the same procedure discussed above to calculate the final p-value in the forward direction *L* → *A* → *B* is also used to calculate a final p-value in the reverse direction *L* → *B* → *A* by swapping the inputs *A* and *B* of NLCD’s tests. Then we use these two p-values, along with a cutoff (significance level of 0.05 say), to make the final call about which model a triplet’s data supports. Specifically as in CIT (Millstein et al., 2009), we call a triplet as:

- causal if forward, but not reverse, p-value is less than 0.05.
- reactive if reverse, but not forward, p-value is less than 0.05.
- independent if both p-values are greater than 0.05.
- inconclusive (no call) if both p-values are less than 0.05.

#### 2.2.6. Optional postprocessing step

Using *M* permutations, the resolution of the empirical p-value of our tests and the smallest achievable empirical p-value are both 1*/*(*M* + 2). For applications requiring higher resolution of small p-values, we can increase *M* ; but relying only on increased permutations has a computational cost. So we employ two steps: increase *M* to a modest 1000 (from the default 500), and postprocess the resulting statistics using a method called EPEPT (Enhanced P-value Estimator for Permutation Tests) (Knijnenburg et al., 2011, 2009). EPEPT employs a generalized Pareto distribution to approximately model the tail of the distribution of permutation-based test statistics. This semi-parametric approach results in higher resolution of small p-values using fewer permutations than the standard non-parametric empirical p-value approach. We downloaded a MATLAB code that implements EPEPT from https://github.com/IlyaLab/epept, and used it to improve the resolution of p-values obtained from applying NLCD to the human GTEx dataset (note that for other datasets considered in this study, the default resolution of p-values was sufficient).

### 2.3. Generation of simulated datasets

To test whether our method’s p-values are sufficiently calibrated to distinguish data from causal vs. independent triplets, we simulated linear and nonlinear datasets pertaining to causal triplets and another dataset pertaining to independent triplets. These datasets were generated under two settings: one where the variances of trait *A* across different strata of *L* are equal, and another where these variances are unequal.

#### 2.3.1. Generation of data with equal variance

Here, the variances of *A* conditioned on different values of *L* are the same. Specifically, we simulated only triplets with haploid genotypes that satisfy:

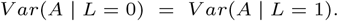

Under this setting, we generated five types of simulated datasets: linear, sine, sawtooth, parabola, and independent. The first four datasets follow the causal model *L* → *A* → *B* with the direct *A* → *B* relation following a linear model for the linear simulated dataset and a nonlinear relation such as sine, sawtooth or parabola for the other datasets. The independent dataset follows the independent *A* ← *L* → *B* model where there is no direct *A* − *B* relation.

To elaborate each dataset further, let’s consider a sine dataset as an example. This dataset comprises 100 different triplets, each of which contains simulated data (*L, A, B* measurements) of *n* individuals/samples. The 100 triplets differ from each other based on their parameter values. There are 10, 5, and 2 distinct values that *µ, σ*_1_, and *σ*_2_ parameters can take respectively, resulting in 10 × 5 × 2 = 100 parameter combinations. For each such combination corresponding to a triplet, its data is generated by independently sampling *n* times from the distributions listed below. The sawtooth dataset is also generated from the same 100 parameter combinations as the sine dataset as shown below. The linear, parabola, or independent dataset is generated using the parameters *β*_1_ and *β*_2_ (with each parameter taking 10 distinct values, resulting in 10 × 10 = 100 parameter combinations or triplets; see below). Data plots of example triplets are in Fig. 2a.

**Fig. 2.**
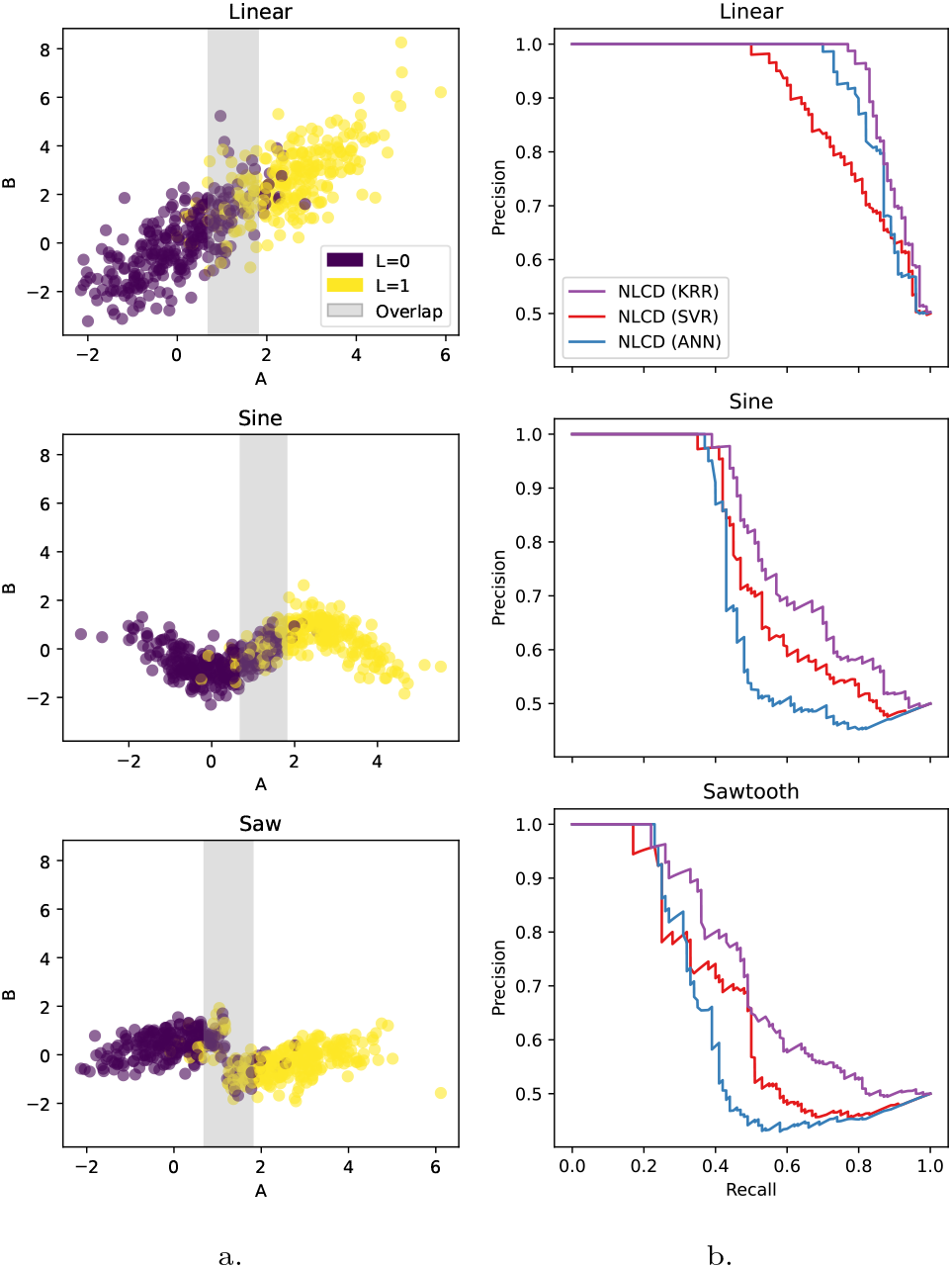
NLCD on simulated benchmark datasets. a. Example plots of the simulated *A* and *B* genes illustrate different types of gene-gene relationships, linear and nonlinear (sine and sawtooth). Shaded grey marks the overlap region between the two gene expression distributions: (*A* | *L* = 0) and (*A* | *L* = 1), with endpoints of this overlap region obtained by setting *P* (*L* = 0 | *A*) to 0.2 and to 0.8 and solving for *A* (see formula derived in Supplementary Section 1.4). b. Performance of NLCD on benchmarks, each comprising data simulated on 100 causal vs. 100 independent triplet models, on a binary (causal vs. independent) classification task is reported as Precision-Recall (PR) curves. Sample size of *n* = 500 is used here.

|  |  |
| --- | --- |
| $L \sim \text{Bernoulli}(0.5)$ | $L \sim \text{Bernoulli}(0.5)$ |
| $\beta_1 \in \{0.1, 0.5, 1, 1.5, 2, 2.5, 3, 4, 5, 6\}$ | $\mu \in \{0.1, 0.5, 1, 1.5, 2, 2.5, 3, 4, 5, 6\}$ |
| $\beta_2 \in \{0.1, 0.5, 1, 1.5, 2, 2.5, 3, 4, 5, 6\}$ | $\sigma_1 \in \{0.2, 0.4, 0.6, 0.8, 1\}$ |
| $A = \beta_1 L + \epsilon_1$ | $\sigma_2 \in \{0.5, 1\}$ |
| $B = \beta_2 f(A) + \epsilon_2$ | $A = \mu L + \epsilon_1$ |
| $\epsilon_1 \sim \mathcal{N}(0, 1) \quad \epsilon_2 \sim \mathcal{N}(0, 1)$ | $B = g\left(\frac{2\pi A}{8\sigma_1} - \frac{\pi}{2}\right) + \epsilon_2$ |
| $f(A) = A \quad (\text{Linear})$ | $\epsilon_1 \sim \mathcal{N}(0, 1) \quad \epsilon_2 \sim \mathcal{N}(0, \sigma_2)$ |
| $f(A) = A^2 \quad (\text{Parabola})$ | $g(\cdot) = \sin(\cdot) \quad (\text{Sine})$ |
| $f(A) = L \quad (\text{Independent})$ | $g(\cdot) = \text{sawtooth}(\cdot) \quad (\text{Sawtooth})$ |

Functions sin(.) and sawtooth(.), from Python’s numpy and scipy libraries respectively, produce the corresponding waveforms with period 2*π*. For instance, sawtooth(.) produces a waveform that increases linearly from −1 to 1 over the interval [0, 2*π*), and jumps back to −1 at the end of each period.

#### 2.3.2. Generation of data with unequal variance

Here, the variances of *A* conditioned on different values of *L* are unequal. Specifically, we simulated only triplets with haploid genotypes that satisfy:

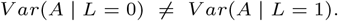

We generated two datasets, linear with unequal trait variance and parabola with unequal trait variance, using the parameter list *β*_1_, *β*_2_, and *α* (each with 5, 5, and 4 distinct values respectively, resulting again in 100 parameter combinations). The simulation process is shown below. Besides simulating triplets with unequal conditional trait variances *β*_1_ ≠ *β*_2_, this process also simulates a smaller fraction of triplets with *β*_1_ = *β*_2_.

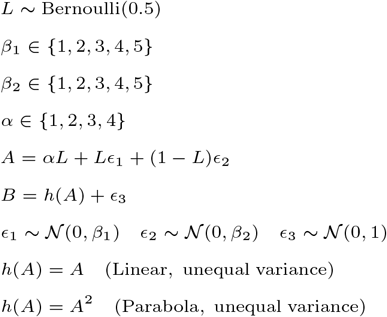

### 2.4. Yeast benchmarking methodology

We evaluated NLCD on a real-world yeast dataset using the benchmarking methodology described in this section.

#### 2.4.1. Yeast data/triplets setup

#### Yeast genomic data and its preprocessing

The yeast genomic data, comprising genotype information, transcriptomic (gene expression) profiles, and covariates of 1012 yeast segregants (derived from a cross of two yeast strains) were obtained from an earlier published study (Albert et al., 2018). Albert et al. had adjusted the gene expression data for the two covariates, experimental batch and a growth-rate covariate, using a linear regression model to remove potential technical confounding effects prior to eQTL mapping. We performed a similar adjustment before further analysis. From the eQTLs detected and reported by Albert et al., we retrieved a list of *cis*-eQTL genes, and the strongest *cis*-eQTL variant for each such gene to select the instrumental variable for the causal discovery methods. For yeast gene annotations, yeast Ensembl release 109 was used.

#### Ground-truth TF → TG data

The ground-truth yeast regulatory relations from transcription factors (TFs) to target genes (TGs) were obtained from the work of (Ludl and Michoel, 2021) (https://github.com/michoel-lab/FindrCausalNetworkInferenceOnYeast), who in turn obtained it from YEASTRACT+ (Teixeira et al., 2022), a curated repository of global transcriptional regulation in yeast species. This repository contains two kinds of experimental evidence for causal TF→TG relations: *DNA binding* evidence (from chromatin immunoprecipitation (ChIP) related experiments) and *expression* evidence (from experimentally-observed expression changes in TGs upon perturbation of a TF). We used ground-truth regulatory relations supported by both DNA binding and gene expression evidence. To derive a subset of these ground-truth relations that are amenable to discovery from Albert et al. data by MR-like causal discovery methods, we retained only: (i) TFs that were identified as *cis*-eQTL genes in the Albert et al. study, and (ii) TGs whose expression was available in Albert et al.’s transcriptomic data. The final ground-truth matrix had 80 TFs along rows, 3422 TGs along columns, and with each matrix entry being either 1 to indicate causal regulation or 0 to indicate non-regulation (of the corresponding TF-TG pair).

#### Assembling the benchmark of yeast triplets

For each TF *A* in the final ground-truth matrix, we assembled triplets of the form (*L, A, B*) by selecting its strongest *cis*-eQTL SNP *L* (i.e., the *cis*-eQTL variant with the highest absolute correlation coefficient to the *A* gene as reported in Albert et al.), and any gene *B* that is associated with *L* at FDR (False Discovery Rate) 20% in Albert et al.’s data. There are 3422 potential *B* genes (TGs or columns of the final ground-truth matrix mentioned above); so to obtain associations at FDR 20%, we test for association between the SNP *L* and each potential *B* gene using Wilcoxon unpaired test, subject the resulting 3422 p-values to Benjamini-Hochberg multiple testing correction, and select *B* genes with corrected p-value ≤ 0.20 (note that this procedure excludes from analysis any self regulation where *B* is the same as *A*). Assembling triplets using this procedure ensures that *L* is a valid instrumental variable associated with both *A* and *B* genes. Note also that this procedure resulted in certain *A* genes with no corresponding *B* genes—such *A* genes are excluded from further analysis.

An assembled triplet (*L, A, B*) is considered (truly) causal if *A* → *B* is 1 in the ground-truth TF → TG matrix, and independent (non-causal) otherwise. For each *cis*-eQTL gene *A*, we found that the independent triplets involving *A* outnumbered the causal triplets involving *A*; so we randomly downsampled the independent triplets to create a balanced dataset with equal number of causal vs. independent triplets for the *A* gene. The resulting *yeast benchmark* contained 1752 causal triplets (involving 66 unique *cis*-eQTL TFs and 1258 unique TGs) and 1752 independent triplets (involving 66 unique *cis*-eQTL TFs and 1395 unique TGs), and is used for all further analysis. Note that each triplet’s data has a sample size of 1012 (due to *L, A, B* being measured across 1012 yeast segregants).

#### 2.4.2. Yeast triplet subsets with different correlation strengths/types

We create different subsets of the triplets in the yeast benchmark by applying cutoffs on measures of linear and nonlinear type correlation, and another measure on the extent of nonlinearity. The resulting high-quality subsets of ground-truth triplets exhibited nonlinearity to varying extents, and can hence be used to find the relative strengths and limitations of different causal discovery methods including ours.

#### Correlation measures used

We employed two correlation measures: Bias Corrected Mutual Information (BCMI) (Pardy, 2013; Pardy et al., 2018) to quantify the strength of any nonlinear association between two variables, and Spearman correlation coefficient (*ρ*) to quantify the strength of linear/monotonic relationship between the variables. In detail, BCMI is a modified (jackknife-based bias-corrected) estimator of mutual information, which in turn is a widely used measure of nonlinearity and non-monotonicity of association between two variables. We preferred Spearman over Pearson correlation (although Pearson directly measures the strength of a linear relationship, and Spearman captures monotonic relationships that include linear associations as a special case) because Spearman is more robust to outliers.

#### Extent of nonlinearity (*δ*)

For each causal triplet in the yeast benchmark, we computed both BCMI and *ρ* between the genes in the triplet using Albert et al. data. We then used the resulting values across all causal yeast triplets to regress |*ρ*| on BCMI, thereby defining an expected relationship between the two measures. We define the extent of nonlinearity (*δ*) of a triplet as the shortest distance of its corresponding point from the regression line, focusing only on points below the regression line to identify nonlinear triplets (i.e., triplets whose |*ρ*| is smaller than expected from the regression, indicating that BCMI is relatively high).

#### Creating correlation-based benchmark subsets

We create different subsets of the yeast benchmark by subjecting its 1752 causal and 1752 independent triplets to different cutoffs (lower bounds) on either BCMI or |*ρ*|, and further subjecting its causal triplets alone to different cutoffs on *δ*. BCMI cutoffs are chosen from 0.05, 0.10, and 0.15; and the corresponding |*ρ*| cutoffs are the predicted |*ρ*| values for each of these BCMI cutoffs using the regression model described above. For each of these cutoff values, we selected triplets with BCMI (or |*ρ*|) at least the cutoff, so that they are sufficiently correlated and hence amenable for causal discovery by the evaluated methods. The result is a high-quality subset of the yeast benchmark, due to removal of several triplets with BCMI (or |*ρ*|) near zero (i.e., triplets whose genes are poorly or not correlated due to noise inherent in real-world data and ground truth not being perfect).

Cutoffs on the extent of nonlinearity, *δ*, are chosen from {0, 0.0125, 0.025, 0.0375, 0.05, 0.1}, such that applying a cutoff of 0 selects all points (causal triplets) below the regression line, 0.0125 selects all points below the regression line at a distance *>* 0.0125 from the line, and so on. These *δ* cutoffs yield different yeast benchmark subsets with varying degree of nonlinearity in the gene-gene association in the causal triplets. We do not subject independent triplets to the *δ* cutoff due to the assumption that the genes in an independent triplet are linked only via *L* and not directly to each other (and hence there is no nonlinear association between the genes on which the *δ* cutoff could be applied).

Just as in the full yeast benchmark, many benchmark subsets created above have more independent than causal triplets. For each such subset, we randomly downsampled the independent triplets to the same number as the causal triplets to ensure class balance (note that other subsets with fewer independent than causal triplets are left untouched). We repeat this process 10 times using different random seeds to obtain 10 different independent triplet lists. Combining each of these independent triplet lists (negative examples) with the causal triplets of the subset (positive examples) yields 10 different random versions of each yeast benchmark subset—performance measures like AUPRC of a benchmark subset are reported using their average and standard deviation across these 10 random versions.

### 2.5. Evaluation of causal discovery methods

We evaluated our NLCD and other existing state-of-the-art (SOTA) methods for causal discovery (CIT, Findr, MRPC) using a classification task and associated benchmark datasets described herein.

#### 2.5.1. Classification task and benchmark datasets

#### Classification task

The binary classification task is to predict whether an input triplet is causal vs. independent. Specifically, the task is to classify whether an input triplet’s data, comprising its (*L, A, B*) measurements across *n* individuals, follows a causal or an independent model. We can evaluate a method performing this task using a benchmark dataset that combines data on a certain number of distinct causal triplets (positive class examples) with that of distinct independent triplets (negative class examples).

#### Simulated benchmark datasets

Each simulated benchmark dataset comprises a simulated dataset on 100 causal triplets and another on 100 independent triplets (see Section 2.3 Generation of simulated datasets). Recall that the linear, sine, or sawtooth simulated dataset comprises data on 100 distinct causal triplets, with each triplet’s data being of sample size *n* and generated using different parameter values of the underlying causal model (see Fig. 2a). Similar to this simulated causal dataset, the simulated independent dataset comprises data on 100 different independent triplets generated using different parameter values of the independent model.

#### Yeast benchmark datasets

The full yeast benchmark dataset contains data on 1752 causal and 1752 independent triplets assembled as described above (see Section 2.4 Yeast benchmarking methodology). Also as described above (see Section 2.4.2 Yeast triplet subsets with different correlation strengths/types), we took nonlinear vs. linear subsets of the causal triplets and matched independent triplets to create different yeast benchmark subsets. We used these benchmark subsets to evaluate NLCD and SOTA methods on triplets with different degrees of nonlinear vs. linear type gene-gene association.

#### 2.5.2. Evaluation measure

AUPRC, calculated using the average precision score, was used to evaluate the performance of the classification models. Though AUPRC can also be calculated using trapezoidal approximation, it involves linear interpolation which is not ideal in the precision-recall space. Average precision score avoids this issue and is also suited for skewed data (Davis and Goadrich, 2006; Flach and Kull, 2015).

#### 2.5.3 Causal discovery calls from evaluated methods

The evaluated methods differ in how they are run on the benchmark triplets’ data (one triplet at a time separately, or on all triplets’ combined data at once), and how they make causality calls (based on applying cutoffs on causality p-values or probabilities). This section provides these details for each evaluated method. Note that the methods are run on each simulated dataset and on the full yeast benchmark dataset. We do not run the methods separately on each yeast benchmark subset; instead, we evaluate performance on these different subsets by extracting and analyzing the relevant causality calls obtained from the full yeast benchmark run.

#### NLCD calls

In our simulated and yeast benchmark datasets, we know the ground truth causal direction. So as discussed in Section 2.2.5 Making the final call, we apply NLCD tests on each triplet only in the forward *L* → *A* → *B* direction; and if the resulting p-value is less than a cutoff (say 0.05), we call the triplet causal, otherwise independent.

#### CIT calls

NLCD is an extension of the CIT framework to capture nonlinear relations, so CIT calls are also made the same way as NLCD calls above.

#### Findr calls

Given a sufficient number of input triplets, Findr (Wang and Michoel, 2017) outputs the probability of a triplet to be causal based on the probability of three of the five component tests implemented in Findr as follows:

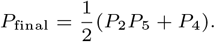

To estimate this probability for each triplet, the findr function `findr.pij_gassist` needs to be applied on the concatenated data of all triplets of interest through a single function call (and not as a separate function call per triplet), as findr needs statistics from a sufficient number of triplets following the null vs. alternate hypotheses to estimate these probabilities. The concatenated data also needs to be organized in a certain fashion as three matrices (Wang and Michoel, 2019) that we explain below, and the above findr function’s output probability matrix needs to be subsetted to extract the causal probabilities of the triplets of interest. More details are in Findr software’s documentation/manual and our code.

To run Findr on each simulated benchmark, we organized the *n* simulated observations on the *m* causal and independent triplets in the benchmark into 3 different matrices: *L*_*mat*_ ∈ {0, 1}^*m*×*n*^, *A*_*mat*_ ∈ ℝ^*m*×*n*^, and *B*_*mat*_ ∈ ℝ^*m*×*n*^. Here, *L*_*mat*_[*i, j*] is the (haploid) genotype of the *i*^*th*^ triplet’s SNP *L* in the *j*^*th*^ individual/sample, *A*_*mat*_[*i, j*] is the expression of the *i*^*th*^ triplet’s *A* gene in the *j*^*th*^ sample, and *B*_*mat*_[*i, j*] is also defined similarly.

To run Findr on the yeast benchmark, we organized the observed genomic data on the causal and independent triplets in the benchmark into three matrices, as done above for the simulated benchmark. In addition, we reordered the rows of these matrices to handle genes that overlap between the sets of *A* and *B* genes across all triplets. Specifically, these overlapping genes were placed first and in the same order in the reordered *A*_*mat*_ and *B*_*mat*_, followed by the other genes (which were also kept in the same order). The *L*_*mat*_ was also reordered accordingly, such that its *i*^*th*^ row corresponds to the genotype values of the strongest *cis*-eQTL/SNP of the *i*^*th*^ gene (row) of *A*_*mat*_.

#### MRPC calls

MRPC was run on each benchmark triplet separately, as this triplet-wise running performed competitively with other methods on the simulated benchmarks, and better than batch-wise running of MRPC on the yeast benchmark as detailed below. We ran MRPC on the data of a given triplet by following these recommendations from its authors: (i) organize data on the triplet into a table with columns in the order *L, A*, and *B*, (ii) compute the robust correlation between all pairs of these three variables using `RobustCor` function of the `MRPC` library with *β* = 0.005 suggested by the authors; and (iii) call the MRPC function in the library with the above data table and robust correlation matrix, and different FDR cutoffs (ranging from 0 to 1, with a step size of 0.05) as its arguments. For each specified FDR cutoff, this function returns the edges among the three nodes of the triplet supported by the data and controlled at this FDR level. Using these edges called at different FDR cutoffs across all the triplets in a benchmark, we can then compute the PR curve of MRPC and associated AUPRC.

For the yeast benchmark, we also tried an alternative batch-wise running of MRPC, wherein MRPC was applied separately to each batch of benchmark triplets sharing the same *A* gene. In detail, for each distinct *A* gene in the benchmark, we selected all triplets involving this *A* gene; organized data on the SNP *L* corresponding to the *A* gene, the *A* gene itself, and all *B* genes in the selected triplets into a data table; computed all pairwise robust correlations among the variables in the table to obtain a correlation matrix; and finally called the MRPC function with arguments: the above data table and correlation matrix, and different FDR cutoffs (ranging from 0 to 1 in steps of 0.05 as in triplet-wise running). The resulting edges among the genes called at different FDR levels were aggregated across all batches to compute the PR curve—the associated AUPRC was poorer than when MRPC was run triplet-wise on the yeast benchmark, and hence we didn’t pursue batch-wise running further.

### 2.6. Application of NLCD to human data: Processing steps

To investigate the potential of NLCD to discover *in vivo* human causal gene networks, we applied NLCD to the GTEx (version 8) human multi-tissue genomic dataset. This dataset comprises transcriptomic measurements from multiple tissue samples obtained postmortem from several hundred human donors, and matched genetic data from the same human individuals. In our applications of NLCD, this human dataset posed challenges due to heterogeneity inherent to a real-world population (i.e., individual-to-individual differences in genetic ancestry, environmental exposures, demographic factors, etc.) and multiple testing burden across the tested triplets. Mitigating these challenges required the selection of tissues with large sample sizes (to tackle reduction of statistical power due to heterogeneity), careful selection of triplets (prior to NLCD’s application to reduce multiple testing burden), and increasing the resolution of p-values (to allow small p-values to survive multiple testing correction), as explained next.

#### Selection of tissues

We selected 10 tissues with the largest sample sizes, and applied NLCD to each of these tissues separately. These top 10 tissues (along with their sample sizes shown within parentheses) are: Skeletal muscle (706), Whole blood (670), Sun-exposed skin (lower leg) (605), Tibial artery (584), Subcutaneous adipose (581), Thyroid (574), Tibial nerve (532), Not sun-exposed skin (suprapubic) (517), Lung (515), and Esophagus mucosa (497). Note that these sample sizes are based on the transcriptomic data available in the GTEx Analysis V8 portal’s open access section named “Single-Tissue *cis*-eQTL Data”, which we refer to as the GTEx data section hereafter.

#### Selection of triplets (for a given tissue)

For any given tissue, we assemble the list of triplets (*L, A, B*) to be tested as follows. We first download from the GTEx data section, the list of *cis*-eQTL relations detected in the tissue at FDR 5% (i.e., q-value cutoff of 0.05 as recommended in the portal). Using the resulting set of *cis*-eQTL SNPs, we next find all significant *trans*-eQTL relations involving any of these SNPs. From these results, we assemble the tissue-specific list of triplets to be tested, by pairing each *cis*-eQTL gene *A* with each *trans*-eQTL gene *B* that are both linked to the same SNP *L*. A *trans*-eQTL relation (*L, B*) is considered significant if its association p-value *<* 10^−5^ and the gene *B* is located at least 1 million base pairs away from *L. Trans*-eQTL relations are detected using the R package Matrix eQTL (Shabalin, 2012), which requires input data on genotype, gene expression, covariates, gene location, and SNP location. Genotype data was extracted from VCF (Variant Call Format) files using VCFTools, following GTEx data-access approval. Other input data such as expression data and covariate data (including genotype principal components and PEER factors) were downloaded from the GTEx data section. Triplets whose eQTL SNP *L* has three distinct genotype values are treated as diploid; those with two distinct values are treated as haploid; and those with one distinct value are excluded from testing.

#### Filtering/Making causal calls (for a given tissue)

For any given tissue, for each selected triplet explained above, we apply NLCD in both directions, *L* → *A* → *B* and *L* → *B* → *A*, to obtain two p-values. Here and in the steps below, *A* refers to the *cis*-eQTL gene and *B* the *trans*-eQTL gene linked to *L*. We explain how these two p-values are used to make causal calls, after certain filters are applied to reduce multiple testing burden and increase the reliability of causal calls.

- We first improved the resolution of NLCD’s four component tests’ p-values and thereby the final p-value using EPEPT as explained in Section 2.2.6 Optional postprocessing step, in order to allow small p-values to remain significant after correction for multiple testing.
- After applying EPEPT, to reduce multiple testing burden and increase confidence in our causal calls, we restricted further analysis to only triplets satisfying three filtering criteria: (i) a Test 3 p-value ≤ 0.1, since testing for causality is unnecessary when *A* and *B* are not correlated; (ii) at least 10 samples for each genotype value; and (iii) high mappability for both genes in the triplet (≥ 0.75 each) and low cross-mappability (0) between them, as sequence similarity between genes can lead to false-positive causal calls (Saha and Battle, 2019). We also tried a relaxed version of the third criteria by additionally including triplets with unknown or unavailable (NA) values for mappability or cross-mappability in the database (Saha and Battle, 2018) used to obtain these measures.
- Consider the triplets retained after the above filtering— their *L* → *A* → *B* test p-values were corrected for multiple testing using the Benjamini-Hochberg method, whereas the reverse *L* → *B* → *A* test p-values were left uncorrected, in order to make more stringent causal calls. Specifically, we call a retained triplet as causal (*L* → *A* → *B*) if its corrected *L* → *A* → *B* p-value is ≤ 0.1 and its (uncorrected) *L* → *B* → *A* p-value is *>* 0.1. Recall that *A* is a *cis*- and *B* a *trans*-eQTL gene linked to *L*; our workflow therefore focuses on making causal calls in the *cis* → *trans* direction, though we acknowledge that *trans*-to-*cis* causal relationships may also exist biologically.

## 3. Results

### 3.1. Our NLCD method overview

NLCD determines if there is a causal relationship between two variables *A* and *B* (e.g., gene expression or other traits) using a third instrumental variable *L* (e.g., a genetic factor correlated to both the traits) by conducting four statistical tests and taking the maximum of these tests’ p-values as the final p-value (see Methods and Fig. 1). Our NLCD’s main contribution is accounting for nonlinear causal relationship between the trait variables in these tests using nonlinear regression (unlike earlier linear causal discovery methods) and employing a novel conditional feature importance score in the fourth test for conditional independence. NLCD can work with any nonlinear regression model, and we have focused on KRR, SVR and ANN in this study (see Methods). The final call on whether data from an input triplet *L, A, B* supports a causal, reactive, or independent model is done based on NLCD’s final p-value as explained in Methods (see 2.2.5 Making the final call).

### 3.2. NLCD outperforms SOTA methods in simulated nonlinear benchmarks

We evaluated our NLCD method’s variants and other SOTA methods on their ability to distinguish causal vs. independent triplets in the simulated benchmark dataset (for sample size *n* = 500 in main text, and other sample sizes in supplement; see also Methods section 2.5.1 Classification task and benchmark datasets). Different types of simulated benchmark datasets correspond to different types of relationship between *A* and *B*, i.e., either a linear relation or a nonlinear relation that takes the form of a sine or sawtooth curve (see Methods section 2.3 Generation of simulated datasets and Fig. 2a).

Among the three variants of our NLCD method applied on different simulated benchmark datasets, we see from Fig. 2b that NLCD using KRR for nonlinear regression gives a better AUPRC than SVR and ANN. We used 500 permutations in these evaluations, and hence we will be using NLCD (KRR) with 500 permutations for further analysis, and refer to it simply as NLCD when it is clear from the context.

We next compared NLCD variants with other SOTA methods for classifying triplets in the simulated benchmark dataset as causal vs. independent, and report the resulting AUPRC and Precision-Recall plots in Fig. 3 and Fig. S1a respectively. We see that NLCD fares well in all the cases, both linear and nonlinear. MRPC does marginally better than NLCD in linear and slightly worse than NLCD in nonlinear. We observe that all the methods perform comparably in the linear benchmarks, and CIT and Findr are worse than other methods in the nonlinear benchmarks. These results were obtained with the number of permutations set at 500 for CIT and NLCD; changing the number of permutations to 100 for both methods didn’t affect the relative performance of CIT and NLCD (see Fig. S1b). As discussed already, the results presented in the main text are on simulated benchmark datasets of sample size *n* = 500. See Table S2 for the AUPRC values of all methods for other sample sizes too.

**Fig. 3.**
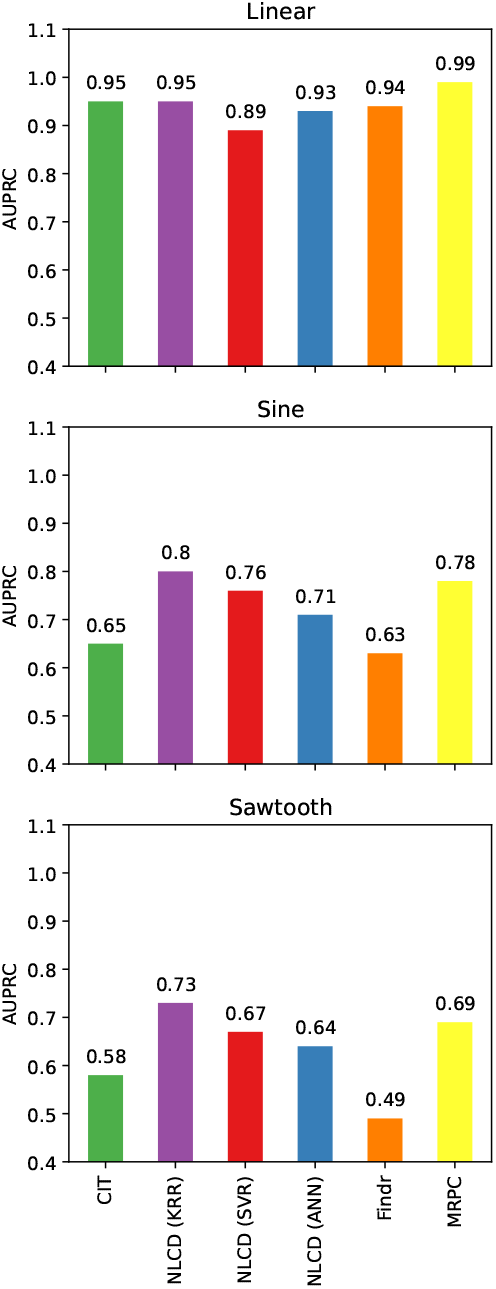
Comparison of NLCD with SOTA methods on simulated benchmark datasets. Simulated benchmarks comprising data on independent triplets vs. causal triplets (with linear, sine, or sawtooth causal relationship) were used to determine the classification performance (AUPRC) of different causal discovery methods (see also Fig. S1a for the underlying PR curves). NLCD and CIT were run using 500 permutations (see also Fig. S1b for results using 100 permutations). Sample size of *n* = 500 is used here.

### 3.3. Detailed evaluation of NLCD relative to the CIT baseline

To complement the comparative evaluations of our NLCD against multiple SOTA methods, we also performed more comprehensive comparison of NLCD against CIT, the baseline method that was extended to develop NLCD, using simulated benchmark datasets.

#### Robustness of performance trends in simulated benchmarks

We first checked whether the better performance of NLCD over CIT on simulated nonlinear (sine or sawtooth) benchmarks persists despite the stochastic nature of both methods. As NLCD and CIT rely on permutation testing, their empirical p-values can change if they are run with different random seeds on the same dataset. To evaluate if the performance trends observed above for one run remain similar across runs, we ran NLCD and the CIT for 10 different runs on the same dataset. The resulting PR curves (see Fig. S2a) show that the performance trends indeed persist despite the methods’ run-to-run variations.

We also checked variability in performance due to variations in the dataset. Ten datasets were simulated using the same generative model (see Methods) but independently using different random seeds. We can see from Fig. S2b that the PR curves of both NLCD and CIT across these 10 datasets show appreciable variability, still NLCD performs better than CIT on nonlinear (sine and sawtooth) benchmarks. The above observations on run-to-run variation and dataset variation hold in the case of both 100 and 500 permutations, and demonstrate that the performance improvement of NLCD method over the CIT baseline on nonlinear benchmarks is robust.

#### Comparing the individual tests of NLCD vs. CIT

To further understand how NLCD extends CIT, we compared the individual tests of these methods. We specifically compared the third test of NLCD and CIT (see Fig. S3), which checks for association between *A* and *B* for each value of *L*. It is observed that across linear triplets, CIT’s and NLCD’s p-values are well correlated; however, across nonlinear (sine or sawtooth) triplets, the p-values of CIT are spread across the y-axis (0 to 1) whereas that of NLCD remain very close to zero. Hence, NLCD works as designed to detect a nonlinear association between *A* and *B* better than CIT. The two methods can also differ in their fourth test, a key test of conditional independence, as follows.

For many nonlinear causal triplets, CIT finds that *L* provides additional information on top of *A* to predict *B*; but this is not the case and *A* alone is sufficient to predict *B* if nonlinearity is properly modeled, which is what NLCD is designed to do and also achieves it via a low CFI score (see also a parabola triplet example below).

#### Performance on unequal variance simulated datasets

Since real-world causal triplets (e.g., yeast genomic data) can exhibit unequal conditional variances of *A* given *L*, and also linear or parabola-like gene-gene relationships, we also tested NLCD and CIT on unequal variance linear/parabola simulated benchmarks. An unequal variance simulated benchmark dataset comprises the simulated dataset on 100 unequal variance linear (or parabola) triplets (see Methods) and another on the 100 equal variance independent triplets. PR curves on this unequal variance benchmark (shown in Fig. S5a for sample size *n* = 500) of NLCD is better than that of CIT, since the former is explicitly designed to handle variance heterogeneity across genotype values. In contrast, both methods perform similarly on equal variance linear and parabola simulated benchmarks (*n* = 500; see Methods and Fig. S5a). Although parabolic relations are nonlinear, they more closely resemble linear associations than sine or sawtooth relations; the linear baseline CIT hence performs comparably to NLCD on the equal variance parabola benchmark. All of these performance trends hold for other sample sizes of 300 and 1000 as well (see Fig. S6).

To illustrate how NLCD handles unequal variances, we simulate an example parabola triplet with unequal conditional trait variances, and plot the predictions of *B* made by NLCD and CIT using *A* and *L* as inputs (see Fig. S5b, S5c, S5d). In the overlap region of these plots, NLCD’s predictions are almost identical between *L* = 0 vs. *L* = 1, whereas CIT’s predictions depend on the value of *L*. Hence for NLCD, *L* doesn’t supplement *A* to predict *B* and thereby results in a low CFI score as mentioned above; whereas for CIT, its linear modeling of a truly nonlinear *A*-*B* relationship results in *L* providing information on top of *A* to predict *B*.

### 3.4. NLCD distinguishes the direction of causality

Benchmark evaluations were done so far in only one direction, i.e., from *A* → *B*. To check if NLCD can infer the direction of causality, simulated data on a benchmark triplet was fed to NLCD separately in the causal direction (*L* → *A* → *B*) and also in the anti-causal direction (*L* → *B* → *A*). Fig. S4 shows that NLCD can distinguish between these directions for causal (linear, sine and sawtooth) triplets, since final p-values output by NLCD are mostly near zero in the causal direction, but spread across the range from 0 to 1 or closer to 1 in the anti-causal direction. For independent triplets which is not causal in either direction, NLCD’s p-values in both directions were near one and thereby supported neither a causal nor reactive model. These observations show that NLCD can correctly identify the causal direction.

### 3.5. NLCD’s performance on the yeast benchmark and its nonlinear subsets

Given the promising performance of our method on simulated benchmark data, we next tested it using real-world yeast genomic data, collected from 1012 yeast segregants. On yeast benchmark datasets comprising causal and independent triplets (see Section 2.4 Yeast benchmarking methodology), we tested whether NLCD and SOTA methods can correctly classify an input triplet as causal vs. independent using genomic data on the triplet. First, note that on the full yeast benchmark dataset, comprising both linear and nonlinear gene-gene associations, Findr and CIT had the highest performance (in terms of AUPRC) followed closely by NLCD and then MRPC (see Fig. S7).

Would the performance trend change if we subset the yeast benchmark to focus on triplets whose genes exhibit sufficient correlation and importantly a high extent of nonlinearity? To answer this question, we assembled benchmark subsets as explained in Methods (Section 2.4.2 Yeast triplet subsets with different correlation strengths/types) by considering the strengths of linear (|*ρ*|) vs. nonlinear (BCMI) type correlation between the genes of triplets (shown in Fig. 4a and Fig. S8 for all causal and all independent triplets respectively in the yeast benchmark). Fig. 4a shows that genes in many causal triplets (i.e., ground-truth causal TF-TG pairs) have low |*ρ*| and low BCMI likely due to noise in genomic data and other factors (see Methods); so it is sensible to filter the benchmark subsets to remove these poorly correlated triplets, which cannot be recovered reliably by any method (including NLCD and CIT as shown in the figure). Another important filter applied on the benchmark subsets retains only triplets with relatively stronger nonlinear than linear type correlation (i.e., more typical BCMI than |*ρ*| values) by applying a lower-bound cutoff on the extent of nonlinearity *δ* (e.g., *δ >* 0.025; see Methods).

**Fig. 4.**
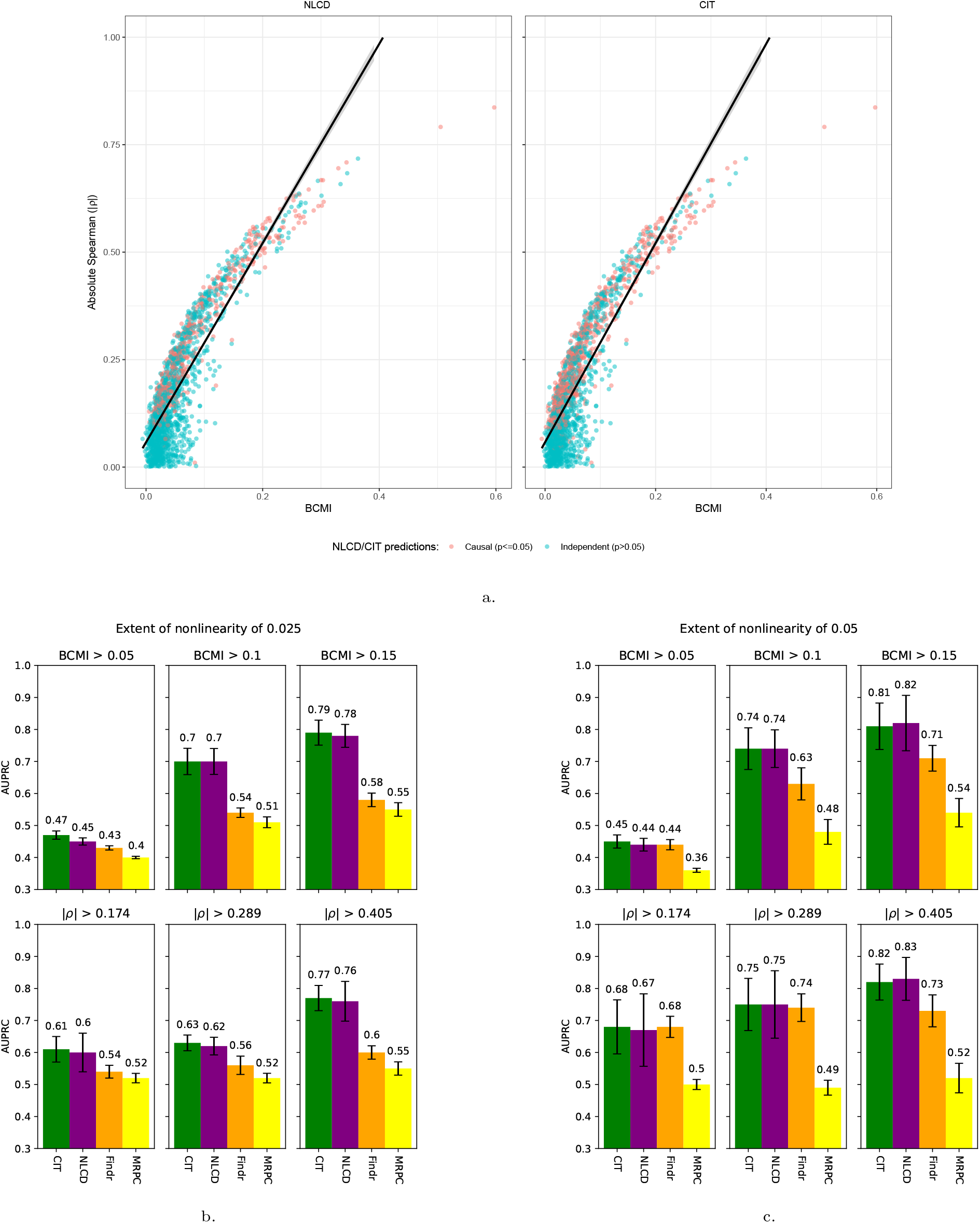
Comparison of NLCD with SOTA methods on the yeast benchmark and its subsets. a. For all causal triplets in the full yeast benchmark (i.e., known TF-TG causal pairs recorded in a ground-truth database), plotted here is the strength of linear (absolute Spearman) vs. nonlinear (BCMI) gene-gene correlation calculated using data from 1012 yeast segregants. Colors indicate triplets predicted as causal vs. independent by NLCD (left) vs. CIT (right) at a (final) p-value cutoff of 0.05. Black line shows the fitted linear regression model. Similar plots are shown for all independent triplets in the full yeast benchmark in Fig. S8. b & c. On different (nonlinear) subsets of the yeast benchmark dataset, performance of causal discovery methods are shown (reported AUPRC is average and standard deviation across 10 random versions of the benchmark subset; random classifier’s AUPRC is 0.5 in all subplots). Here, we show the plots for an extent of nonlinearity (*δ*) of at least 0.025 and 0.05 (see Fig. S9 for other cutoff values).

Across several subsets of the yeast benchmark dataset with different degrees of nonlinearity, our NLCD and the CIT method performed comparably/better than other methods (see Fig. 4b and Fig. 4c, and similar figures for more *δ* cutoffs in Fig. S9). In detail, for benchmark subsets compiled using low BCMI cutoff (e.g., BCMI *>* 0.05), AUPRC of all the methods are mostly worse than the random AUPRC of 0.50 for reasons already explained above pertaining to insufficiently correlated causal gene pairs. However at higher cutoffs on BCMI and across different cutoffs on *δ*, AUPRC improves appreciably for all the methods, with CIT and NLCD in particular showing consistently better performance than Findr. MRPC’s performance was lower and may be attributed to how it was run on the triplets (however note that we tried different ways of running MRPC on the yeast benchmark and report the best-performing one here; see Methods). NLCD is comparable to but not strictly better than CIT for the *δ* cutoffs tested, and may perform better in more nonlinear benchmark subsets where the *δ* cutoff is increased beyond 0.05. But such subsets could not be assembled due to the relatively small number of nonlinear triplets in the yeast benchmark. In summary, taking all the results together, NLCD is a competitive method for recovering causal triplets from this real-world yeast genomic data.

### 3.6. Discovering causal gene relations across multiple human tissues

The promise of NLCD is to detect causal relations under *in vivo* human settings, where interventions are not possible and observational data is the main resort to discover causality. But heterogeneity in a human population that leads to weak instruments and multiple testing burden across all tested triplets poses challenges, and necessitates large sample sizes and careful selection of triplets. So, we applied NLCD to only the 10 human tissues with the most number of samples in GTEx and filtered the triplets using *cis*/*trans*-eQTL associations (see Methods 2.6 Application of NLCD to human data: Processing steps). Furthermore, we increased resolution of the NLCD tests’ p-values by using 1000 instead of the default 500 permutations and postprocessing the resultant test statistics using the EPEPT method (see Methods 2.2.6 Optional postprocessing step).

We first ran our NLCD workflow on the skeletal muscle tissue triplets, which had a sample size of 706, and summarize the results in Fig. 5. Specifically, we plot in Fig. 5a the p-values from test 3 (of conditional association) and test 4 (of conditional independence) for these muscle triplets, as these test 3 and 4 p-values are indicative of correlation and causation respectively. This plot shows that only a subset of the correlated gene pairs show strong evidence of causality, thereby emphasizing the importance and challenge of distinguishing causality from correlation in human datasets of moderate sample sizes. At stringent p-value and mappability/cross-mappability thresholds (see Methods), we discovered two causal triplets (Fig. 5f). One of them is the *IRF1-PSME1* pair – it is known from literature (Shi et al., 2011) that *IRF* 1 is a transcription factor regulating a target gene *PSME*1. When inspecting the data and NLCD predictions for this regulating pair (see Fig. 5b and 5c), we see that the NLCD predictions for all three values of *L* are close to each other in the overlap region (where the IRF1 values corresponding to these three *L* values overlap), thereby leading to the critical conditional independence test 4 passing. To provide a counterexample, we show similar plots (Fig. 5d and Fig. 5e) for a pair (*IRF1-PAP10*) that is not causal but correlated, to indicate how the NLCD predictions for different values of *L* are different in the overlap region.

**Fig. 5.**
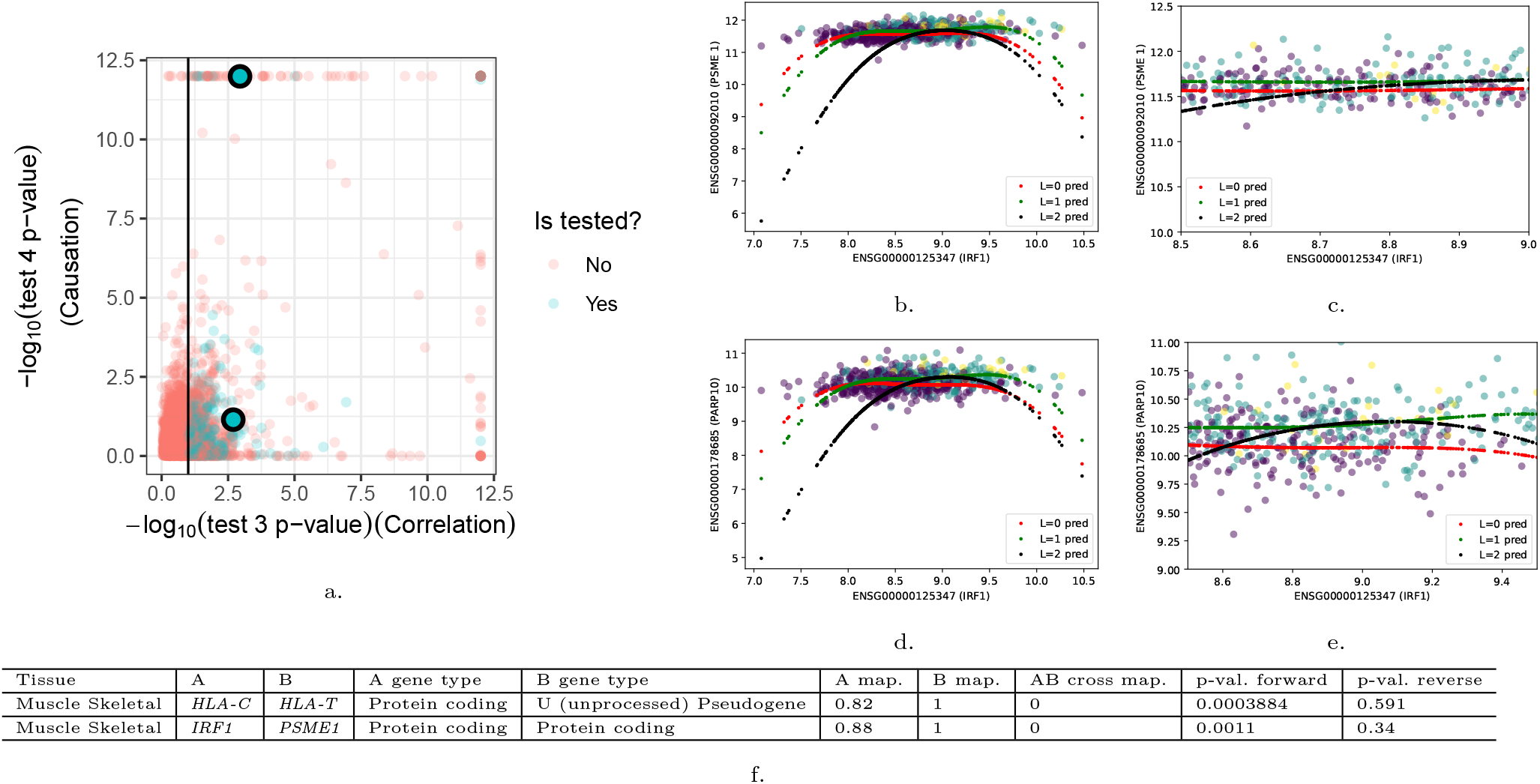
Application of NLCD to human muscle GTEx data. a. Strength of causal vs. correlative relationship of the gene pair in each muscle triplet, as measured via the corresponding NLCD tests’ p-values, is shown. Only triplets satisfying certain filtering criteria (see Methods 2.6 Application of NLCD to human data: Processing steps) were tested; and colored as indicated in the legend. The black line with x-intercept of 1 corresponds to a p-value cutoff of 0.1. The two black-circles indicate the gene pairs *IRF1*-*PMSE1* and *IRF1*-*PAP10*. b & c. NLCD’s predictions (denoted pred) of *PMSE1* using *IRF1* and different values of *L* as inputs is plotted (left), along with a zoomed view of this plot (right). d & e. NLCD’s predictions (denoted pred) of *PAP10* using *IRF1* and different values of *L* as inputs is plotted (left), along with a zoomed view of this plot (right). f. Listed are all muscle triplets detected as causal by NLCD based on stringent p-value (pval.) and mappability/cross-mappability (map./cross map.) thresholds.

We next ran NLCD on all the 10 selected GTEx tissues, after relaxing the mappability/cross-mappability criteria (See Methods section: 2.6 Application of NLCD to human data: Processing steps). Note that with the original stringent criteria, NLCD could not pick up causal triplets in any non-muscle tissue – this could be due to lower sample sizes in non-muscle compared to muscle tissue. The relaxed criteria resulted in 14 causal triplets (Table 1). Most of the discovered pairs, involving pseudogenes, cannot be dismissed as mappability-related artifacts, as the role of pseudogenes as genuine regulators has been suggested earlier (Pink et al., 2011). Our predicted causal relations, both on pseudo genes and protein-coding genes, provide promising hypotheses that could be tested further using other datasets and follow-up experiments.

**Table 1.**
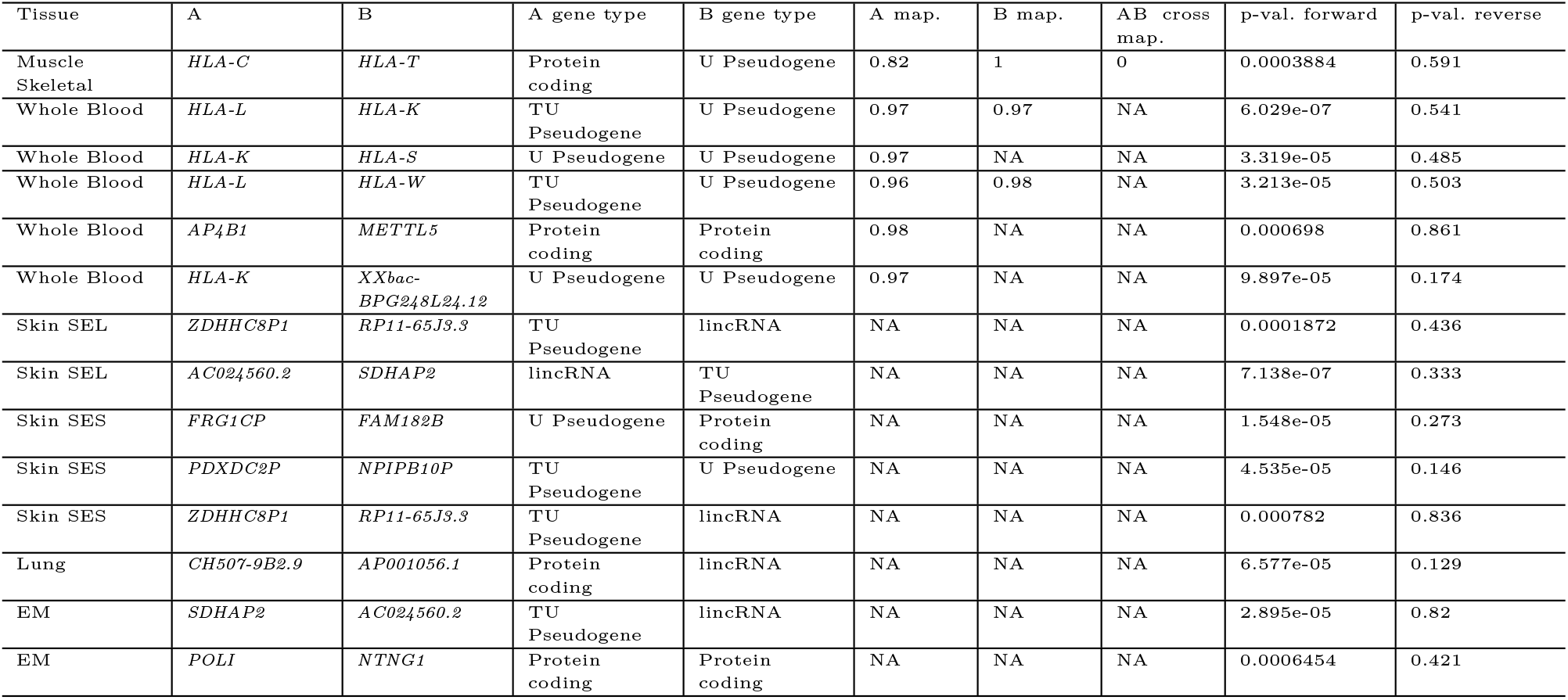
Human causal triplets detected by NLCD under relaxed criteria. Causal triplets detected by NLCD in the selected human GTEx tissues are listed as in Fig. 5f, but after relaxing a filtering criterion: triplets with unknown (NA) mappability/cross-mappability values for their genes are not removed. All other filtering criteria and significance cutoffs remain the same as before (see Methods 2.6 Application of NLCD to human data: Processing steps). Abbreviations: Skin Sun Exposed Lower Leg (Skin SEL), Skin Not Sun Exposed Suprapubic (Skin SES), Esophagus-Mucosa (EM), Transcribed Unprocessed (TU), Unprocessed (U), lincRNA (long intergenic non-coding RNA).

## 4. Conclusion and Discussion

We’ve developed a method NLCD to account for nonlinearity when discovering causal relationship between two gene expression or clinical traits. NLCD works under an MR-like framework wherein a genetic factor associated with the two tested traits is used to infer the presence of causality from observational data on this triplet. The evidence for causal relationship between the traits/genes in the triplet is summarized using a p-value based on four statistical tests.

Existing SOTA methods for the triplet-based causal discovery problem assume linear relationship between the genes. NLCD, by virtue of its NLR modeling and CFI scoring to realize the statistical tests, shows promising performance on nonlinear simulated and real-world benchmarks. Specifically, in simulated benchmark datasets comprising triplets with nonlinear gene-gene causal relationships, NLCD achieves better AUPRC than existing SOTA methods. For triplets with linearly related genes, NLCD performs comparably to other methods. In real-world yeast benchmarks, NLCD is competitive with other methods in recovering triplets with linear or nonlinear TF-TG causal relationships. Application of NLCD to every pair of genes with a common eQTL SNP provides a workflow to uncover the gene regulatory network underyling a biological system, and we illustrate this approach on human multi-tissue genomic data to detect tissue-specific causal genes.

Some caveats about NLCD’s performance are worth mentioning. NLCD is developed by extending a linear causal discovery method CIT. Whereas NLCD clearly outperformed the linear baseline CIT in simulated nonlinear benchmarks, NLCD and CIT performed similarly on real-world yeast triplets. This may be due to the predominantly linear causal relationship between TFs and TGs in the yeast genomic dataset. NLCD’s advantage over CIT is likely to become more pronounced in practice, if the real-world datasets involve complex clinical traits, unequal conditional trait variances, or time-series measurements of periodic genes. Application to human GTEx data showed that NLCD is likely to be useful in practice for detecting reliable causal relations only when sample sizes of cohorts are sufficiently large (close to 1000s) to overcome weak genetic instruments and multiple testing burden, and stringent filters are applied to reduce false positives driven by (cross-)mappability issues. NLCD was able to overcome these issues to a reasonable extent and thereby detect reliable causal relations using yeast segregants’ and human GTEx data; larger sample sizes of observational data and longitudinal measurements in future studies such as biobank-scale human studies can make it easier for NLCD to discover causal networks.

We motivate certain future directions to improve NLCD methodology further. The running time of NLCD (KRR) increases cubically with sample size and linearly with the number of permutations (see Table S3 and Table S4). We could incorporate faster KRR methods (Alaoui and Mahoney, 2015) into NLCD as it could help us run more permutations in a given time period and thus obtain higher resolution p-values. NLCD’s current default is to use 500 permutations, the minimum number recommended in an earlier study (Marozzi, 2004). Another future work pertaining to NLCD methodology would be to conduct its tests 3 and 4 (i.e., compute the respective test statistics of MSE loss and CFI score) using a testing dataset. Currently, the whole dataset on a triplet *D*_*T*_ is used to train the model as well as calculate the test statistics. In the future, we could assess whether the already competitive performance of NLCD on benchmarks can be further improved by splitting *D*_*T*_ into a training and a testing dataset for model fitting and test statistics computation respectively. Theoretical studies on the sample complexity of NLCD could be another direction of future work. Specifically, given our generative model of data from a causal triplet, can we derive a lower bound on the number of samples *n* required to detect this causal triplet with high probability? This lower bound could be a function of the strength of the causal relation, the size of (i.e., the fraction of *n* datapoints that lie in) the overlap region between different values of *L*, and other relevant factors. Finally, while we have focused on CIT as the baseline to extend upon for nonlinear causal discovery, we can also develop a nonlinear extension of any other triplet-based causal discovery method by incorporating the core concepts of NLCD (viz., NLR modeling and CFI scoring) in their methodology.

NLCD makes a minimal set of assumptions and simple choices for the NLR models to determine the presence of causality. To assess whether NLCD is robust against deviations from its simple distributional assumptions, we inspected NLCD’s behavior on simulated datasets whose conditional distributions deviated from normality (see Supplementary Section 2.1 NLCD’s tests under departures from normality, and Table S5 and Table S6). These results showed that NLCD can recover causal triplets even when the genes do not follow a normal distribution. However, we note that NLCD’s reliance on permutation testing implies that the different steps of NLCD are modular and therefore its distributional assumptions can be changed in a fairly straightforward manner if required. NLCD is also model-agnostic with respect to NLR, and nonlinearity can be modeled using any other suitable model besides the three used here.

To conclude, we have developed a method NLCD for observational-data-driven linear and nonlinear causal discovery, and demonstrated its competitiveness in diverse datasets. These results hold promise for its future applications to rapidly accumulating data on genetic/molecular/clinical variables from large (e.g., biobank-scale) studies to reveal causal biological networks operating *in vivo*.

## 5. Acknowledgements

We would like to thank Neeraj Rajkumar Parmaar for his contribution during the initial phase of this project. We would like to thank Rahul Biswas, Brintha V P, Nilesh Subramanian, Ritwiz Kamal and all the other BIRDS (Bioinformatics and Integrative Data Science) lab members for their feedback on this work, and their support. We would like to thank Harish G Ramaswamy, Debarghya Ghoshdastidar and Arun Rajkumar for their valuable inputs on the conditional independence test and related aspects of this work. We acknowledge the use of LLMs (Large Language Models, mainly ChatGPT, Gemini, and Grammarly) for copy-editing purposes (checking spelling and grammar, and polishing certain sentences/phrases).

## Data Availability

The code implementing our method NLCD and its application to simulated or real-world datasets can be found here: https://github.com/BIRDSgroup/NLCD.

