## Supplementary Information for "NLCD: A method to discover nonlinear causal relations among genes"

(for the manuscript "NLCD: A method to discover nonlinear causal relations among genes" by Aravind Easwar and Manikandan Narayanan)

### 1. Supplementary Methods

#### 1.1. Background on underlying theorem and model

NLCD is based on the underlying causal equivalence theorem from an earlier study (Chen et al., 2007):
Given a triplet $(L, A, B)$ with the assumption that $L$ is "randomized", there exist a causal relation $L \to A \to B$ and no confounders exist that are causal for both $A$ and $B$ if and only if the following conditions hold:

- $L \sim A$
- $L \sim B$
- $L \perp B \mid A$

In the above theorem, we say that $L$ is "randomized" if each individual's $L$ is set randomly to one of its values (not necessarily uniformly at random), before and without regard to, i.e., independent of, the individual's $A$ and $B$ values. Each individual's $L, A, B$ values are also independent of each other and identically distributed. These are standard assumptions in the MR framework and are supported by random recombination and assortment that happen during reproductive cell formation and fertilization.

Based on this theorem, a method can be developed to infer causality by implementing the tests for association between $L$ and $A$, and $L$ and $B$, and the key test for conditional independence between $L$ and $B$ given $A$. These tests, along with an additional conditional independence test of $L$ and $A$ given $B$, can help discriminate between the three models, causal, reactive and independent, mentioned in the main text.

#### 1.2. Specifications of the nonlinear models used

We performed nonlinear regression (NLR) using the following models: Artificial Neural Network (ANN), Support Vector Regression (SVR) and Kernel Ridge regression (KRR). We called the functions implementing these NLR models in the scikit-learn package with the hyperparameter values shown below (all other function arguments were set to default values):

ANN specifications:

- activation function: ReLU where $\text{ReLU}(x) = \max(0, x)$
- solver (for weight optimization): Adam optimizer
- alpha (regularization parameter): 0.0001
- max iterations: 200
- hidden layer: one with 100 neurons

SVR specifications:

- kernel: rbf (radial basis function)
- C (regularization parameter): 1
- epsilon (of $\epsilon$-SVR model): 0.1
- gamma: $\frac{1}{(\text{number of features}) * \text{variance}(X)}$, where $X$ is the data matrix containing the values of all features across all samples, and $X$ is viewed as a vector containing all these values for the purpose of computing the variance here

KRR specifications:

- kernel: rbf
- alpha (regularization parameter): 1.0

#### 1.3. Calculation of probabilities for CFI score

At first, the chosen model is trained with $L$ and $A$ as inputs, and $B$ as the output. Then the prediction is made for $A$ where $L = 0$ and for $A$ where $L = 1$. We take the difference in predictions in the cases above where there is an overlap of the predictions. Using Bayes' theorem, the probability is calculated. This is done by calculating the probability of the regions where the data is present. The Bayes formula applied here is

$$p(L = 0 \mid A_t) = \frac{p(A_t \mid L = 0)\ p(L = 0)}{p(A_t)} \tag{1}$$

Similarly, for the other term, $p(L = 1 \mid A_t)$.

The term $p(A_t \mid L = 0)$ is found from the normal distribution whose mean is given by the sample mean of all the points of $A$ where the points are $L = 0$, and the standard deviation of the distribution is given by the sample standard deviation of these points. Similarly $p(A_t \mid L = 1)$ is calculated. The term $p(L = 0)$ is estimated as the ratio of samples having $L = 0$ by the total number of samples, similarly for $p(L = 1)$. The normalizing factor $P(A_t)$ is given by

$$P(A_t) = P(A_t \mid L = 0)\ P(L = 0) \quad + \quad P(A_t \mid L = 1)\ P(L = 1) \tag{2}$$

#### 1.4. Overlap region of triplet gene distributions conditioned on genotype

For a triplet visualized in Fig. 2a in the main text, we shade in grey a region where the gene expression distributions under different values of SNP, i.e., $(A \mid L = 0)$ and $(A \mid L = 1)$, overlap. The endpoints of the overlap region were obtained by setting $P(L = 0 \mid A) = 0.2$ and $P(L = 0 \mid A) = 0.8$ and solving them using the formula derived here.

Specifically, we seek a formula to derive the value $x$ of $A$ where $P(L = 0 \mid A = x) = p$ for some value $p$. If $p$ is set to 0.2 and 0.8, then the corresponding values of $A$ form an interval that can be considered as the overlap region where the distributions $(A \mid L = 0)$ and $(A \mid L = 1)$ overlap.

We have

$$P(L = 0 \mid A) = \frac{P(A \mid L = 0)P(L = 0)}{P(A \mid L = 0)P(L = 0) + P(A \mid L = 1)P(L = 1)}$$

$$P(L = 0 \mid A) = \frac{1}{1 + \frac{P(A|L=1)P(L=1)}{P(A|L=0)P(L=0)}}$$

Say $P(L = 0)$, the fraction of samples where $L = 0$ is $a$, then $P(L = 1) = 1 - a$. Say $P(L = 0 \mid A) = p$, rearranging the terms we have,

$$p = \frac{1}{1 + \frac{P(A|L=1)\cdot(1-a)}{P(A|L=0)\cdot a}}$$

$$\frac{P(A \mid L = 1) \cdot (1 - a)}{P(A \mid L = 0) \cdot a} = \frac{1 - p}{p}$$

Taking $\log_e$ on both sides,

$$\log_e \frac{1 - a}{a} + \log_e P(A \mid L = 1) - \log_e P(A \mid L = 0) = \log_e \frac{1 - p}{p}$$

Since we assume that the conditional distribution of $A$ given $L = \ell$ is normal with mean $\mu_\ell$ and variance $\sigma_\ell^2$, we've:

$$P(A = x \mid L = 0) = \frac{1}{\sqrt{2\pi\sigma_0^2}} e^{-\frac{(x-\mu_0)^2}{2\sigma_0^2}}, \text{ and}$$

$$P(A = x \mid L = 1) = \frac{1}{\sqrt{2\pi\sigma_1^2}} e^{-\frac{(x-\mu_1)^2}{2\sigma_1^2}}.$$

Substituting these values, and using $\text{logit}(a) = \log_e \frac{1 - a}{a}$, we get:

$$\text{logit}(a) + \left[-\frac{(x - \mu_1)^2}{2\sigma_1^2}\right] + \left[\frac{(x - \mu_0)^2}{2\sigma_0^2}\right] - \log_e \sigma_1 + \log_e \sigma_0 = \text{logit}(p)$$

$$\text{logit}(a) - \frac{(x^2 + \mu_1^2 - 2x\mu_1)}{2\sigma_1^2} + \frac{(x^2 + \mu_0^2 - 2x\mu_0)}{2\sigma_0^2} - \log_e \sigma_1 + \log_e \sigma_0 = \text{logit}(p)$$

Rewriting,

$$\left(\frac{1}{2\sigma_0^2} - \frac{1}{2\sigma_1^2}\right) x^2 + \left(\frac{\mu_1}{\sigma_1^2} - \frac{\mu_0}{\sigma_0^2}\right) x + \left(\frac{\mu_0^2}{2\sigma_0^2} - \frac{\mu_1^2}{2\sigma_1^2} - \log_e \sigma_1 + \log_e \sigma_0 + \text{logit}(a) - \text{logit}(p)\right) = 0$$

This is of the form,

$$r \cdot x^2 + q \cdot x + s = 0$$

$$\text{So,} \quad x = \frac{-q \pm \sqrt{q^2 - 4rs}}{2r}.$$

We find the $x$ value satisfying $P(L = 0 \mid A = x) = p = 0.2$ and another $x$ value satisfying $P(L = 0 \mid A = x) = p = 0.8$, and use these $x$ values as the endpoints of the desired overlap region (note that the corresponding values of $P(L = 1 \mid A = x)$ are 0.8 and 0.2 respectively).

### 2. Supplementary Results

#### 2.1. NLCD's tests under departures from normality

We wanted to check if NLCD was robust against departures from its model assumptions – specifically, we wanted to check the performance of NLCD on triplets where the gene's conditional distributions deviate from normality. To do so, we considered our simulated datasets (linear, sine, sawtooth and parabola) and segregated the simulated genes into genes that followed a normal distribution conditioned on $L$ and those that did not. We employed the Lilliefors test, which is based on the Kolmogorov–Smirnov test, for checking normality (Lilliefors, 1967). In detail, we checked each $A$ gene in the simulated dataset for normality as follows: take the minimum of p-values given by the Lilliefors test for $A \mid L = 0$ and $A \mid L = 1$ distributions as the final Lilliefors p-value; and if this p-value is less than 0.05, call the gene as following a non-normal distribution (see Table S5). We also performed a similar check for each $B$ gene in the simulated data (see Table S6). From the two tables, we can see that NLCD is capable of detecting causality in triplets that exhibit a non-normal behavior. This is especially true in linear and parabolic datasets. In sine and sawtooth datasets, we see that there are non-normal triplets that are not called causal, however that is also the case for normally distributed triplets, suggesting that these wrong calls may be due more to insufficient sample sizes than deviations from normality.

### 3. Supplementary Tables

| L | A | B |
|---|---|---|
| 0 | 0.46 | -0.34 |
| 0 | 0.24 | 1.031 |
| 0 | 0.90 | -0.63 |
| 1 | 0.00 | 0.66 |
| 0 | 0.50 | 1.33 |
| 1 | 1.30 | 2.12 |
| 0 | 0.86 | 1.90 |
| 0 | -2.23 | 5.47 |

**Table S1. Example dataset of a triplet.** For a triplet $(L, A, B)$, this table shows an example dataset with each row being an observation of the triplet. Each observation comprises values of the genotype variable $L$ and trait variables $A$ and $B$ in a sampled individual.

| Simulated benchmark dataset sample sizes | NLCD | | | CIT | | | Findr | | | MRPC | | |
|---|---|---|---|---|---|---|---|---|---|---|---|---|
| | Linear | Sine | Sawtooth | Linear | Sine | Sawtooth | Linear | Sine | Sawtooth | Linear | Sine | Sawtooth |
| 300 | 0.94 | **0.77** | 0.68 | 0.92 | 0.62 | 0.52 | 0.93 | 0.55 | 0.48 | **0.97** | 0.72 | **0.68** |
| 500 | 0.95 | **0.80** | **0.73** | 0.95 | 0.65 | 0.58 | 0.94 | 0.63 | 0.50 | **0.99** | 0.78 | 0.69 |
| 1000 | 0.94 | **0.79** | 0.76 | 0.96 | 0.64 | 0.60 | 0.94 | 0.66 | 0.56 | **1.00** | 0.76 | **0.77** |

**Table S2. Performance of causal discovery methods on simulated benchmark datasets across different sample sizes.** AUPRC values of methods, NLCD, CIT, Findr, and MRPC, for classifying the triplets in different simulated benchmark datasets of varying sample sizes as causal vs. independent. Shown in bold are the best performing method's AUPRC for each sample size and type of relation (linear, sine and sawtooth) of the causal triplets in the benchmark.

| Permutations | Sample size | Running time (seconds) |
|---|---|---|
| 100 | 300 | 4.74 |
| | 500 | 12.41 |
| | 1000 | 72.47 |
| 500 | 300 | 21.68 |
| | 500 | 55.15 |
| | 1000 | 355.03 |

**Table S3. Running time of NLCD.** Running time taken by NLCD for different number of permutations and sample sizes. NLCD was run on Supermicro SSG-6039P-E1CR16H server with 112 cores, each containing an Intel(R) Xeon(R) Platinum 8180 2.50GHz CPU; and a total RAM of 1TB. Parallelization of triplets' processing was achieved by using the multiprocessing library of Python.

| Tests | Time Complexity | Remarks |
|---|---|---|
| Test 1 | $O(nM)$ | Computation of NLL takes $O(n)$ and it is done $M$ times plus one time for the original data |
| Test 2 | $O(n^3) + O(nM)$ | For regressing out, we are using KRR, which takes $O(n^3)$. Again calculating the NLL score $M$ times similar to Test 1. |
| Test 3 | $O(n^3M)$ | For each value of genotype, we are permuting $B$ and fitting the model using KRR. |
| Test 4 | $O(n^3M)$ | For each permutation, we calculate the CFI score that requires fitting the model using KRR. |
| **Overall** | $O(n^3M)$ | |

**Table S4. Runtime complexity of NLCD.** Time complexity of the four statistical tests in NLCD in terms of $n$, the number of samples, and $M$, the number of permutations.

| Simulated Dataset | NLCD | Lilliefors p-value | |
|---|---|---|---|
| | | $< 0.05$ | $\geq 0.05$ |
| Linear | $p < 0.05$ | 5 | 72 |
| | $p \geq 0.05$ | 0 | 23 |
| Sine | $p < 0.05$ | 1 | 35 |
| | $p \geq 0.05$ | 6 | 58 |
| Sawtooth | $p < 0.05$ | 1 | 20 |
| | $p \geq 0.05$ | 7 | 72 |
| Parabola | $p < 0.05$ | 4 | 76 |
| | $p \geq 0.05$ | 1 | 19 |

**Table S5. NLCD results stratified by the normality of $A$.** Number of triplets split according to the non-normality of $A$ gene and the causal calls made by NLCD. Lilliefors test p-value of less than 0.05 suggests that the distribution is not normal. These are the same (equal variance) datasets underlying the simulated benchmarks studied in the main text, with sample size $n = 500$; and NLCD was run using 100 permutations.

| Simulated Dataset | NLCD | Lilliefors p-value | |
|---|---|---|---|
| | | $< 0.05$ | $\geq 0.05$ |
| Linear | $p < 0.05$ | 4 | 73 |
| | $p \geq 0.05$ | 1 | 22 |
| Sine | $p < 0.05$ | 15 | 21 |
| | $p \geq 0.05$ | 27 | 37 |
| Sawtooth | $p < 0.05$ | 14 | 7 |
| | $p \geq 0.05$ | 14 | 65 |
| Parabola | $p < 0.05$ | 5 | 75 |
| | $p \geq 0.05$ | 0 | 20 |

**Table S6. NLCD results stratified by the normality of $B$.** Number of triplets split according to the non-normality of $B$ gene and the causal calls made by NLCD. Lilliefors test p-value of less than 0.05 suggests that the distribution is not normal. These are the same (equal variance) datasets underlying the simulated benchmarks studied in the main text, with sample size $n = 500$; and NLCD was run using 100 permutations.

### 4. Supplementary Figures

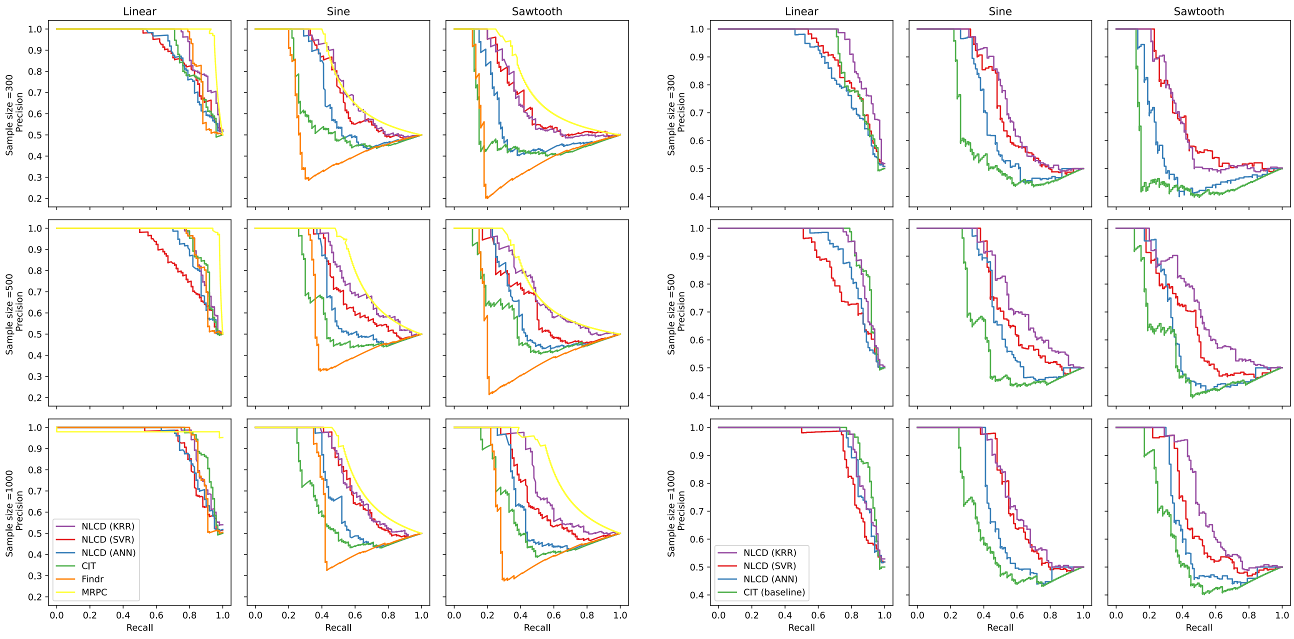


(a) (b)

**Fig. S1. PR curves of causal discovery methods applied on simulated benchmark datasets.** a. PR curve of different methods for sample sizes 300, 500, and 1000, with the number of permutations for CIT and NLCD set at 500. b. PR curve of NLCD and CIT, in particular, with the number of permutations set at 100. Note that the y-axis range of panel b is different from that in panel a to zoom into the plots.

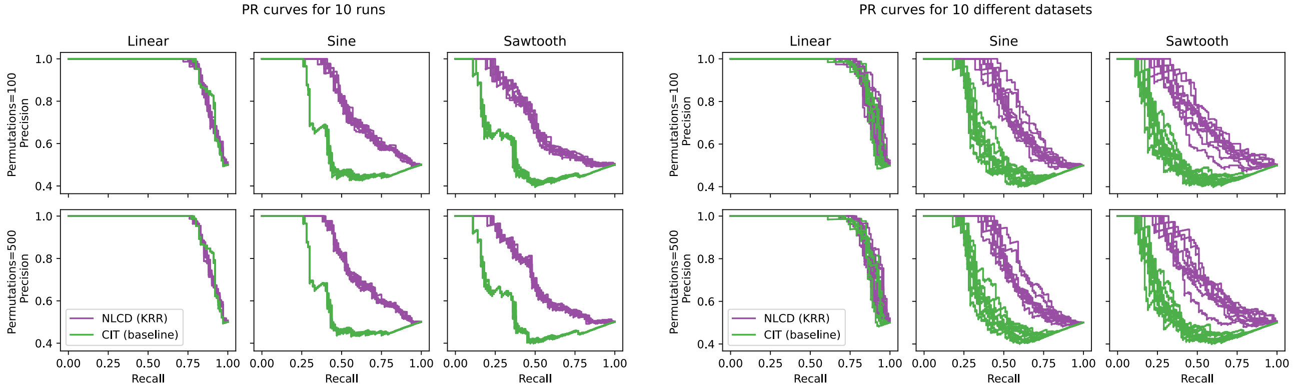


(a) (b)

**Fig. S2. Performance of CIT/NLCD under run-to-run and dataset-to-dataset variations.** a. For each type of simulated benchmark, a single benchmark dataset is simulated, and the PR curves of 10 runs of NLCD and of CIT on this single dataset are shown. b. For each type of simulated benchmark, 10 benchmark datasets are simulated (using different random seeds), and the PR curves of a single run of NLCD and of CIT on these 10 datasets are shown. All simulated benchmark datasets used here have sample size $n = 500$, and NLCD and CIT were run using either 100 or 500 permutations as indicated in the figure.

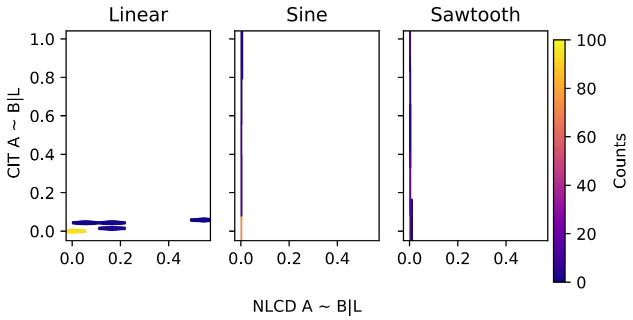


**Fig. S3. Comparison of "Test 3" p-values of CIT/NLCD.** The hexagonal bin plots compare p-values of the conditional association test (Test 3) of CIT and its corresponding test in NLCD across the causal triplets of different simulated benchmark datasets (with sample size $n = 500$).

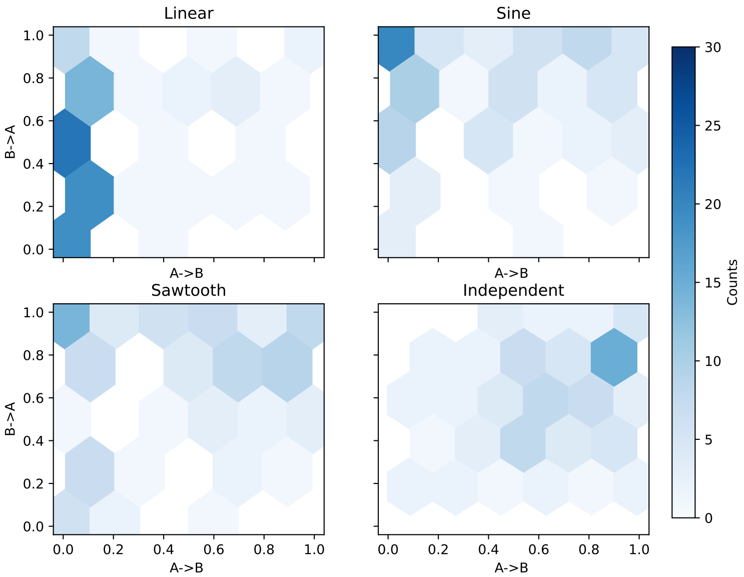


**Fig. S4. Bidirectional application of NLCD.** Comparison of p-values of NLCD when applied in $A \to B$ or $B \to A$ direction on simulated datasets that are generated in the A→B direction. These are the same datasets underlying the simulated benchmarks studied in the main text, with sample size $n = 500$.

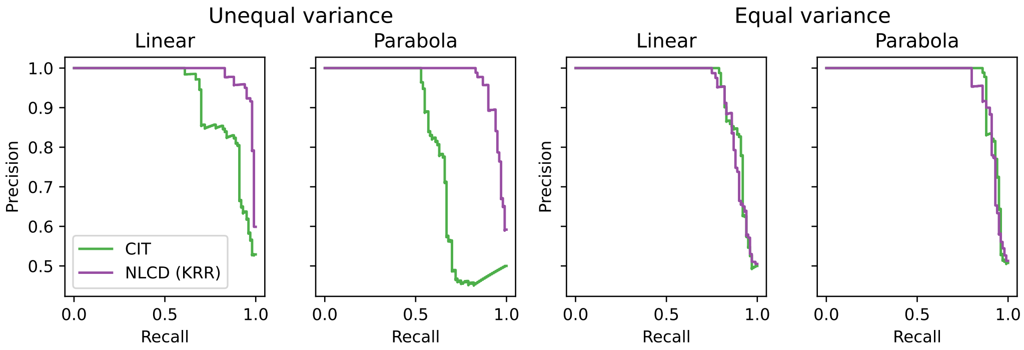


(a)

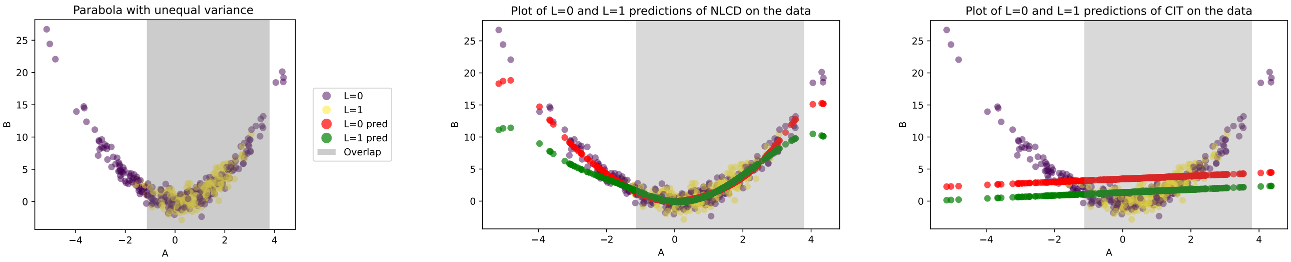


(b) (c) (d)

**Fig. S5. Effect of unequal variance on CIT vs. NLCD, and closer inspection of a parabola causal triplet.** a. Shown are the PR curves of CIT and NLCD run using 100 permutations on linear and parabola simulated benchmarks with sample size $n = 500$, and conditional variances of $A$ being unequal (first two plots) or equal (next two plots) given different values of $L$. b. Plot of an example triplet, specifically a simulated unequal variance causal parabola triplet. c. NLCD's prediction of $B$ from $A$ and $L$ for the unequal variance parabola triplet in panel b. d. CIT's prediction of $B$ from $A$ and $L$ for the unequal variance parabola triplet in panel b. In panels c and d, the term "pred" stands for prediction, and red dots indicate the $B$ predictions using $L = 0$ and the corresponding $A$ values, and green dots the $B$ predictions using $L = 1$ and the corresponding $A$.

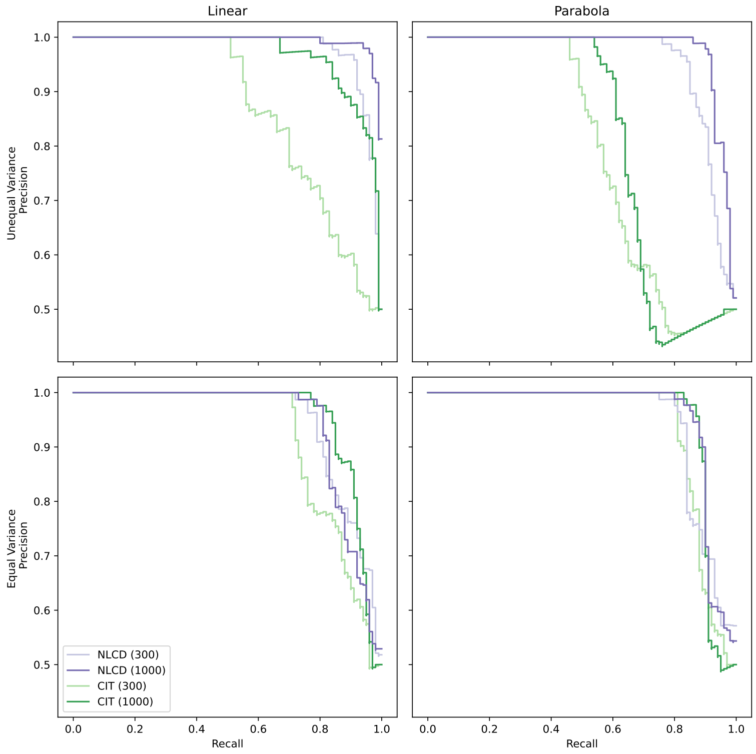


**Fig. S6. Effect of unequal variance on CIT and NLCD under different sample sizes.** Performance of CIT and NLCD on linear and parabola simulated benchmark datasets, when the variances of the $A$ gene conditioned on different genotype ($L$) values are equal (bottom) vs. unequal (top). The benchmark datasets had a sample size of 300 or 1000 as indicated within parentheses in the legend; and the methods were run using 100 permutations.

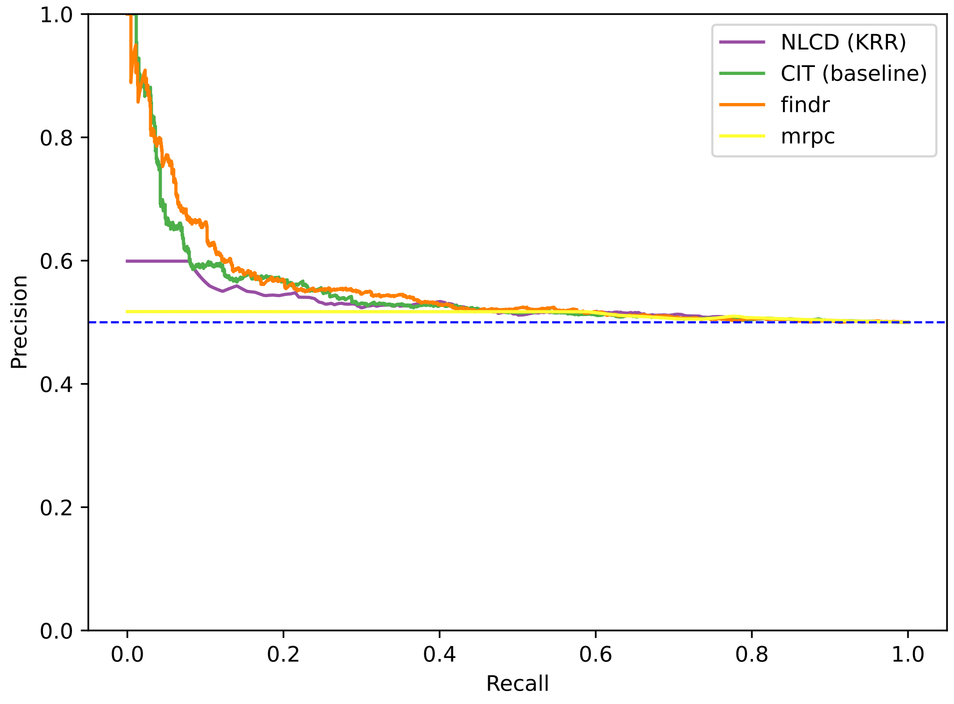


**Fig. S7. PR curves on the full yeast benchmark.** PR curves of NLCD and SOTA methods on the task of classifying triplets in the full yeast benchmark dataset as causal vs. independent. The full benchmark includes triplets with linear and/or nonlinear type gene-gene correlations. The curves are plotted using hyperbolic interpolation (Davis and Goadrich, 2006), and the corresponding AUPRC values are: 0.55 (Findr), 0.55 (CIT), 0.53 (NLCD), and 0.51 (MRPC). Random classifier's AUPRC is 0.50.

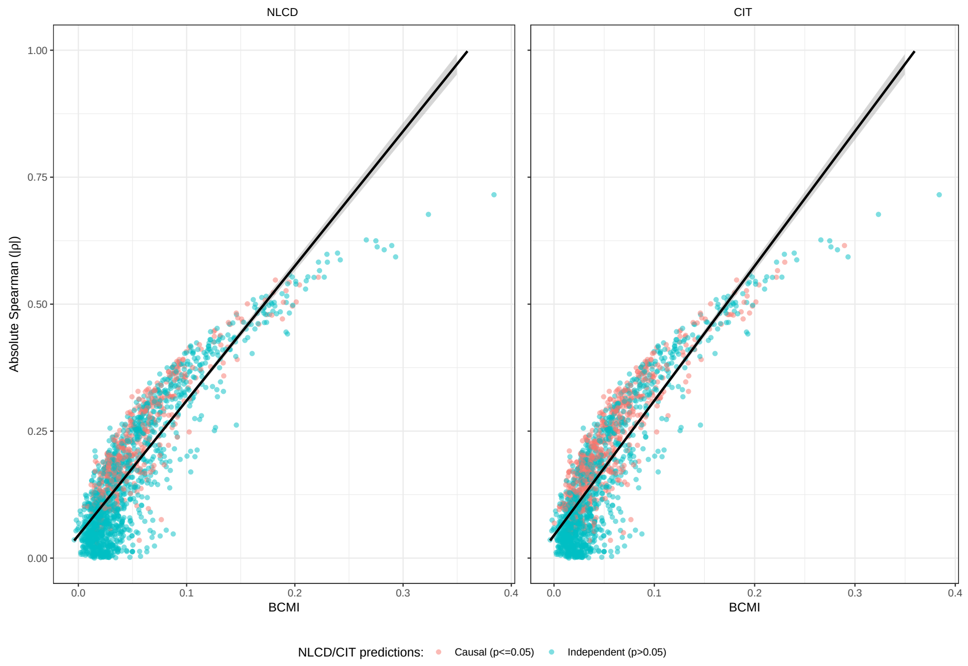


**Fig. S8. NLCD and CIT predictions on the yeast benchmark's independent triplets and all triplets.** For all independent triplets in the full yeast benchmark (i.e., TF-TG pairs recorded as independent/non-causal in a ground-truth database; see Methods in main text), plotted here is the strength of linear (absolute Spearman) vs. nonlinear (BCMI) gene-gene correlation calculated using data from 1012 yeast segregants. Colors indicate triplets predicted as causal vs. independent by NLCD (left) vs. CIT (right) at a (final) p-value cutoff of 0.05. Black line shows the fitted linear regression model.

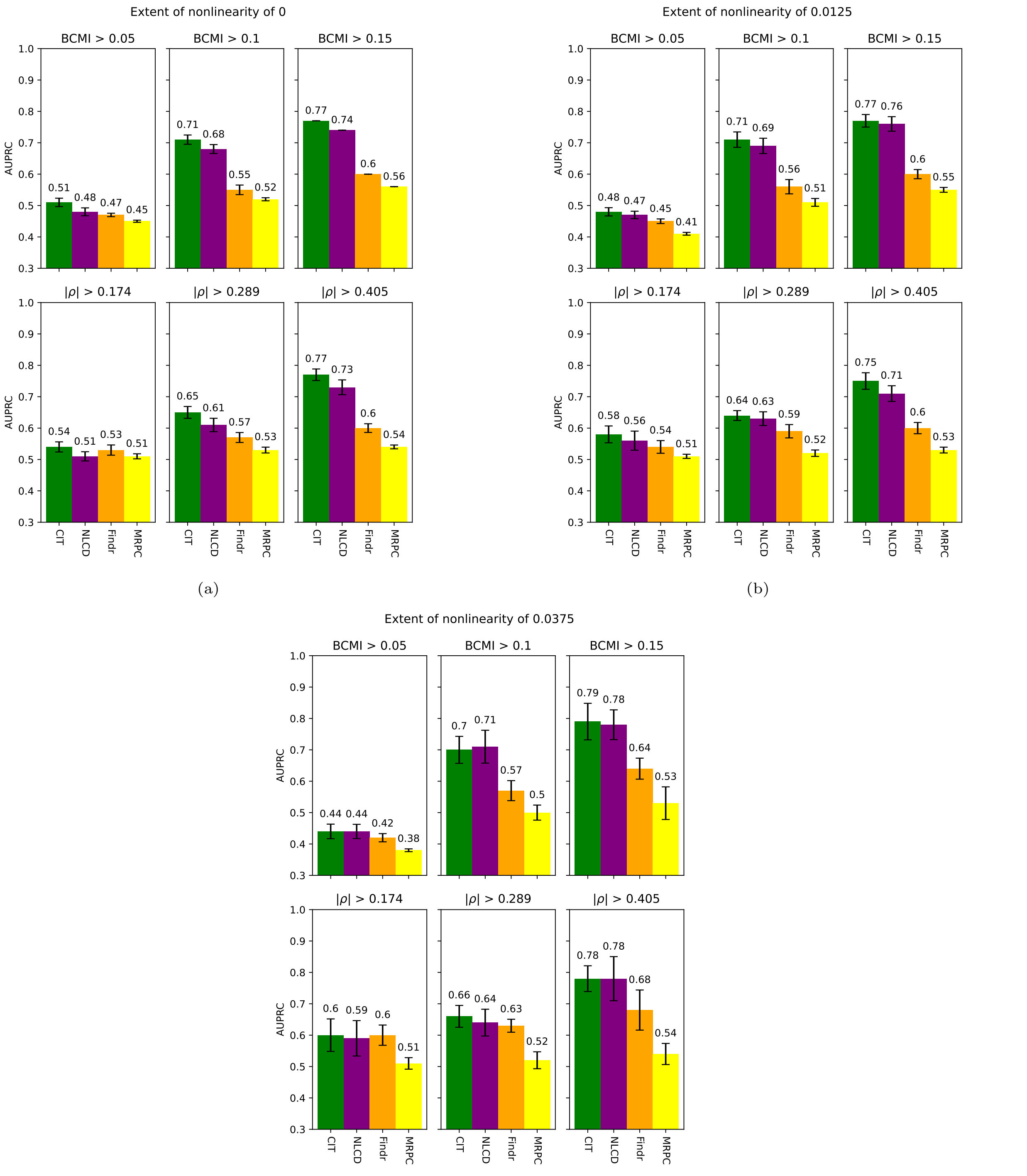


(c)

**Fig. S9. Comparison of NLCD with SOTA methods on further (nonlinear) subsets of the yeast benchmark.** On different (nonlinear) subsets of the yeast benchmark dataset, performance of causal discovery methods are shown (reported AUPRC is average and standard deviation across 10 random versions of the benchmark subset; random classifier's AUPRC is 0.5 in all subplots). Here, we show the plots for an extent of nonlinearity ($\delta$) of at least 0, 0.0125, and 0.0375, to complement similar plots shown in Fig. 4b and Fig. 4c in the main text for other $\delta$ cutoffs.
